# Active mechanics of sea star oocytes

**DOI:** 10.1101/2022.04.22.489189

**Authors:** Peter J. Foster, Alexandra Zampetaki, Jinghui Liu, Sebastian Fürthauer, Nikta Fakhri

**Affiliations:** Physics of Living Systems, Department of Physics, Massachusetts Institute of Technology, Cambridge, MA, 02139, USA; Department of Physics, Brandeis University, Waltham, MA; Department of Physics and Astronomy, Bridge Institute, Michelson Center for Convergent Bioscience, University of Southern California, Los Angeles, CA, 90089, USA; Institute for Applied Physics, TU Wien, A-1040 Wien, Austria; Max Planck Institute of Molecular Cell Biology and Genetics, Dresden, SN, 01307, Germany; Center for Computational Biology, Flatiron Institute, New York, NY, 10010, USA

## Abstract

Cell shape changes, driven by the contractile actomyosin cortex, are essential for multicellular organisms. Yet, how contractility and cell shape changes emerge from molecular scale interactions between proteins in the actomyosin cortex remains unclear. Oocytes of the bat star *Patiria miniata* exhibit a traveling wave of cellular deformation during meiosis, known as a surface contraction wave (SCW). Here, we exploit this highly stereotypical deformation to study how cortical properties are set by microscale processes. By pharmacologically modulating the levels of polymerized actin, we show that the wildtype composition maximizes the cellular deformation rate. To understand these results, we developed an active fluid model for shape changes that demonstrates that deformation rates are set by the ratio between cortical viscosity and active contractile stresses and provides a framework for deriving these two key material properties from coarse-grained molecular-scale interactions. In this model, the ratio of active stress to viscosity peaks at intermediate admixtures of passive crosslinkers. This explains the observations of the drug treatment experiments and makes additional predictions, namely that overexpression of either myosin or passive crosslinkers would decrease deformation rates, which we verified experimentally. Together, this shows that the interplay between cortical active stress and viscosity relies on balancing the ratio of myosin to passive crosslinkers on each filament, which can be modulated by actin density. Our results argue that changing the relative molecular compositions of cortical crosslinks and motors provides a mechanism that biological systems could exploit for robust and predictable control over both viscosity and active stresses.

## Introduction

Actomyosin networks are canonical examples of living matter. They generate nonequilibrium active stresses by converting chemical energy from their environment into mechanical work [1–5]. In cells, actomyosin is the major component of the cellular cortex, which is the prime actuator of cell shape changes. Here, we use a naturally occurring surface contraction wave (SCW) in meiotic *Patiria miniata* oocytes as a platform for studying the mechanisms by which actomyosin-driven cell shape changes are enabled.

The mechanisms underlying the contractility of disordered actomyosin networks such as the cortex remain poorly understood [6–8]. Contractility in disordered actomyosin networks has been shown to depend not solely on myosin activity, but on the the architecture [8, 9] and density [10, 11] of the actin network. Furthermore, a substantial body of work in *in vitro* systems has demonstrated that in many cases F-actin and myosin alone are insufficient for network contractility and that additional actin crosslinking proteins are required [12–16], though an exception has been found at low pH where myosin itself can function as an effective crosslinker [17].

Understanding how the cellular-scale properties of actomyosin networks emerge from the filament-scale interactions of the network’s constituents is an open challenge. To generate contractile stresses, the filament-scale symmetry between contraction and expansion must be broken [18]. A number of microscopic models have been proposed for how this symmetry breaking can occur [8, 19]. One class of models relies on polarity sorting - myosin accumulates at actin barbed ends, clustering barbed ends together which in turn leads to isotropic contraction [20]. Myosin-2 end accumulation has been demonstrated in a purified system [7], and this mechanism has been argued to give rise to contraction in microtubule networks [21–23]. Alternatively, contractility has been proposed to arise from the nonlinear mechanical properties of F-actin, which can buckle under compression. In purified systems, F-actin buckling has been seen to coincide with network contraction, [16, 24]. Finally, contractility independent of myosin motor activity has been proposed for some structures, such as the contractile F-actin shell that captures chromosomes during sea star oocyte meiosis [25–29]. However, directly assessing the degree of myosin end accumulation or filament buckling *in vivo* presents an experimental challenge due to the high density and small size of myosin and actin filaments, which limits the ability to resolve individual motors and filaments using light microscopy.

Here, we consider the actomyosin-driven surface contraction wave of meiotic *Patiria miniata* oocytes as a model for cellular contractility. Using pharmacological inhibitions targeting actin polymerization dynamics, we find that cellular deformation during the contraction wave is not a monotonic function of cortical actin density, but is instead peaked near the wild type density. To understand the physical origin of this observation, we first developed a physical theory for deforming active surfaces, which shows that the rate of deformation is proportional to the ratio between the active contractile cortical stress and the cortical viscosity. This led us to conclude that our pharmacological treatments directly affect this key parameter. We note that Rho GTPase signaling, which has been shown to be a key determinant of surface contraction wave speed and spatial patterning [30, 31], provides the upstream biochemical input that generates the traveling band of active stress in our system and remains largely unaffected between experiments. Our mechanical model takes this traveling active stress as given and asks how the cortical material properties set the resulting cellular deformation. The two descriptions are thus complementary: signaling determines where and when the cortex contracts, while mechanics determines how much the oocyte deforms. We next sought to understand how controlling actin density can affect the ratio between the active contractile cortical stress and the cortical viscosity.

To relate the values of this key ratio to the molecular composition of the cortex, we utilized a recently developed theoretical framework for dense cytoskeletal networks [32], which generalizes a model developed for microtubule networks [33] to allow for more elaborate motor and crosslinker properties. Based on this, we developed an active fluid model coarse-grained from a microscopic description of actin, crosslinkers, and motors. This model makes quantitative predictions for how the rate of oocyte deformation varies with the concentrations of passive active crosslinkers and active motor proteins, namely that the radial deformation rate slightly increases before decreasing as passive crosslinker concentration increases, and surprisingly, decreases monotonically with increasing active motor concentration. We compared these predictions with experimental measurements from oocytes overexpressing *α*-actinin or myosin regulatory light chain and found quantitative agreement. Taken together, these results provide a conceptual bridge from filament-level microscopic interactions, to cellular scale material properties of the cell cortex, and ultimately to the emergent mechanics that enables whole-cell deformations *in vivo*.

While the result that deformation rates scale with the active stress-to-viscosity ratio is consistent with established active gel theory [3], our contributions are: (i) the first *in vivo* demonstration that this ratio controls a natural stereotyped deformation, (ii) explicit closed-form expressions for viscosity and active stress from a microscopic motor-crosslinker model, enabling quantitative predictions, and (iii) experimental verification of the counterintuitive prediction that myosin overexpression decreases the deformation rate.

## Results

### Surface contraction wave dynamics

As a model process for actomyosin-driven contraction *in vivo*, we here consider the surface contraction wave preceding the first meiotic division in oocytes of the bat star *Patiria miniata* [31, 34, 35]. First discovered in developing axolotl [36], surface contraction waves are found in a variety of large eggs including those of the frog *Xenopus laevis* [37], barnacles [38], and ascidians [39]. In sea star oocytes, these waves are driven by a band of activated Rho that travels across the oocyte from the vegetal to animal pole, guided by a spatial gradient of cdk1-cyclinB [31, 35, 40]. This traveling band of active Rho locally activates several downstream factors, including myosin via the ROCK pathway and the actin nucleator formin mDia1 [41]. These in turn lead to local contraction, resulting in a traveling surface contraction wave (SCW), whose arrival at the animal pole coincides with polar body extrusion [31, 34](Fig. 1a, Supplementary Video 1). Due to the ease of meiotic induction, the large, spherical shape of sea star oocytes, and the highly conserved nature of the actomyosin components [18], sea star oocytes are a powerful model system for the study of actomyosin contractility *in vivo*. While a surface contraction wave coincides with each meiotic division, we here consider the first surface contraction wave which takes place during meiosis i. Previous pioneering work[31] has uncovered several key determinants of surface contraction waves in this system. Here, we build on this foundation and develop a quantitative framework for bridging from protein-scale interactions to cellular-scale deformation in this system.

**Figure 1:**
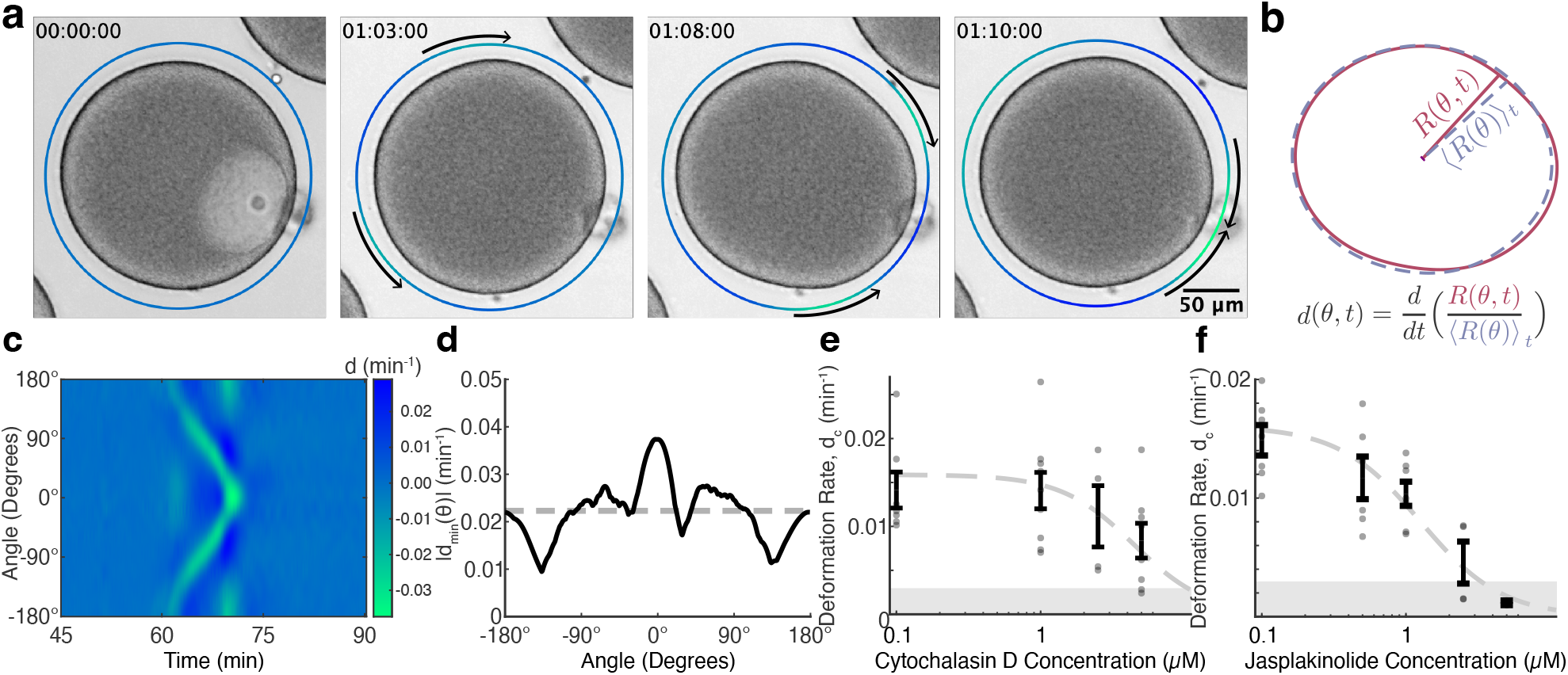
Surface contraction wave dynamics. (a) Timelapse of oocyte surface contraction wave. The outer circle color denotes *d*(*θ, t*), the radial deformation rate for each angle, colored as in c. Negative (green) values indicate contraction while positive (dark blue) values indicate expansion. Arrows indicate the direction of wave propagation. (b) Deformation rate calculation. For each angle at each time, the radial distance between the oocyte’s centroid and outer contour was normalized by the time averaged radial distance before a time derivative was taken. (c) Kymograph of *d*(*θ, t*). The surface contraction wave is readily visualized as the converging lines of negative (green) values indicating contraction. (d) Solid line: Magnitude of the minimum deformation rate for each angle, *d*_*min*_(*θ*). Dashed line: *d*_*c*_, the average value of |*d*_*min*_(*θ*)|. (e, f) Average characteristic deformation rate, *d*_*c*_, as a function of cytochalasin D (e) and jasplakinolide (f). Gray dots: measurements from individual oocytes. Black lines: average deformation rate ± s.e.m. Gray regions: noise floor, defined as the average deformation rate measured for non-maturing, germinal vesicle stage oocytes (n=9). Dashed line: IC50 fits. Cytochalasin D experiments: n=4 to n=9 oocytes per treatment condition. Data was collected over 9 days using oocytes from 6 clutches. Jasplakinolide experiments: n=2 to n=7 oocytes per treatment condition. Data was collected over 6 days using oocytes from 4 clutches.

We first quantified a characteristic radial deformation rate and the wave propagation speed during the SCW. To quantify the deformation rate, the distance between the oocyte’s center and outer contour at each angle and time point, *R*(*θ, t*) was first normalized by the time averaged radial distance for that angle, ⟨*R*(*θ*)⟩_*t*_ and a time derivative was taken to compute the local deformation rate, *d*(*θ, t*) (Fig. 1b, see *Materials and Methods*). From kymographs of *d*(*θ, t*), the SCW can be readily visualized as a traveling line of negative values (Fig. 1c), and the propagation speed of the wave can be measured from the slope of this line (see *Materials and Methods*). A characteristic deformation rate was calculated by first taking the magnitude of the minimum deformation rate for each angle, |*d*_*min*_(*θ*)| (Fig. 1d, solid line) which was then averaged across angles to determine the characteristic deformation rate, *d*_*c*_ = ⟨|*d*_*min*_(*θ*)| ⟩_*θ*_ (Fig. 1d, dashed line, see *Materials and Methods*). Control oocytes were found to have a mean wave propagation speed of *v* = 47 ± 4 *µm/min* (mean ± s.e.m., n=25 oocytes), consistent with previous measurements [31], and a characteristic deformation rate of *d*_*c*_ = 0.017 ± 0.002 min ^−1^ (mean ± s.e.m., n=25 oocytes).

### Characteristic deformation rate is maximum at intermediate cortical actin density

We next investigated how modulating actin density through perturbing actin turnover influences deformation dynamics during the SCW. We first considered the actin polymerization inhibitor cytochalasin D, which at high concentrations has been shown to inhibit deformation during the SCW [31]. Measurements of the characteristic deformation rate, *d*_*c*_, and the wave speed, *v*, were repeated for oocytes treated with varying concentrations of cytochalasin D. As expected for an actomyosin-driven process, the characteristic deformation rate was found to monotonically decrease with increasing cytochalasin D concentration (Fig. 1e). Fitting a dose response curve yielded an IC_50_ of 4.4 ± 2.0 *µ*M (fit value ± 95% confidence interval, see *Materials and Methods*). For concentrations of cytochalasin D ≤ 5 *µ*M, where deformation during the SCW was large enough for the wave speed to be measured, no significant differences in wave speed were found between treatment conditions (Extended Data Figure 1), consistent with previous results arguing that the speed of the SCW is set by the spatiotemporal dynamics of cdk1-cyclinB [31, 35].

We next considered the effects of jasplakinolide, which induces actin polymerization and stabilization [42]. We found a dose-dependent decrease in the characteristic deformation rate (Fig. 1f), consistent with previous results where actin stabilization with phalloidin suppresses the deformation wave [31]. Fitting a dose response curve yielded an IC_50_ of 1.3 ± 0.8*µ*M (fit value ± 95% confidence interval, see *Materials and Methods*). Thus, treatments with drugs that alter actin polymerization reveal that the characteristic deformation rate is maximized at intermediate actin densities.

To quantitatively assess how these perturbations modulate actin density, we next overexpressed and imaged LifeAct-mCherry and used its fluorescence intensity as a proxy for F-actin density (see *Materials and Methods*). As expected, LifeAct-mCherry localized to the oocyte’s periphery and to the nuclear region shortly after the onset of nuclear envelope breakdown, (Fig. 2a), consistent with F-actin’s role in nuclear envelope breakdown in sea star oocytes [43, 44]. As time progresses, the cortical LifeAct-mCherry signal globally decreases before locally increasing slightly during the surface contraction wave (Fig. 2a, Supplementary Video 2).

**Figure 2:**
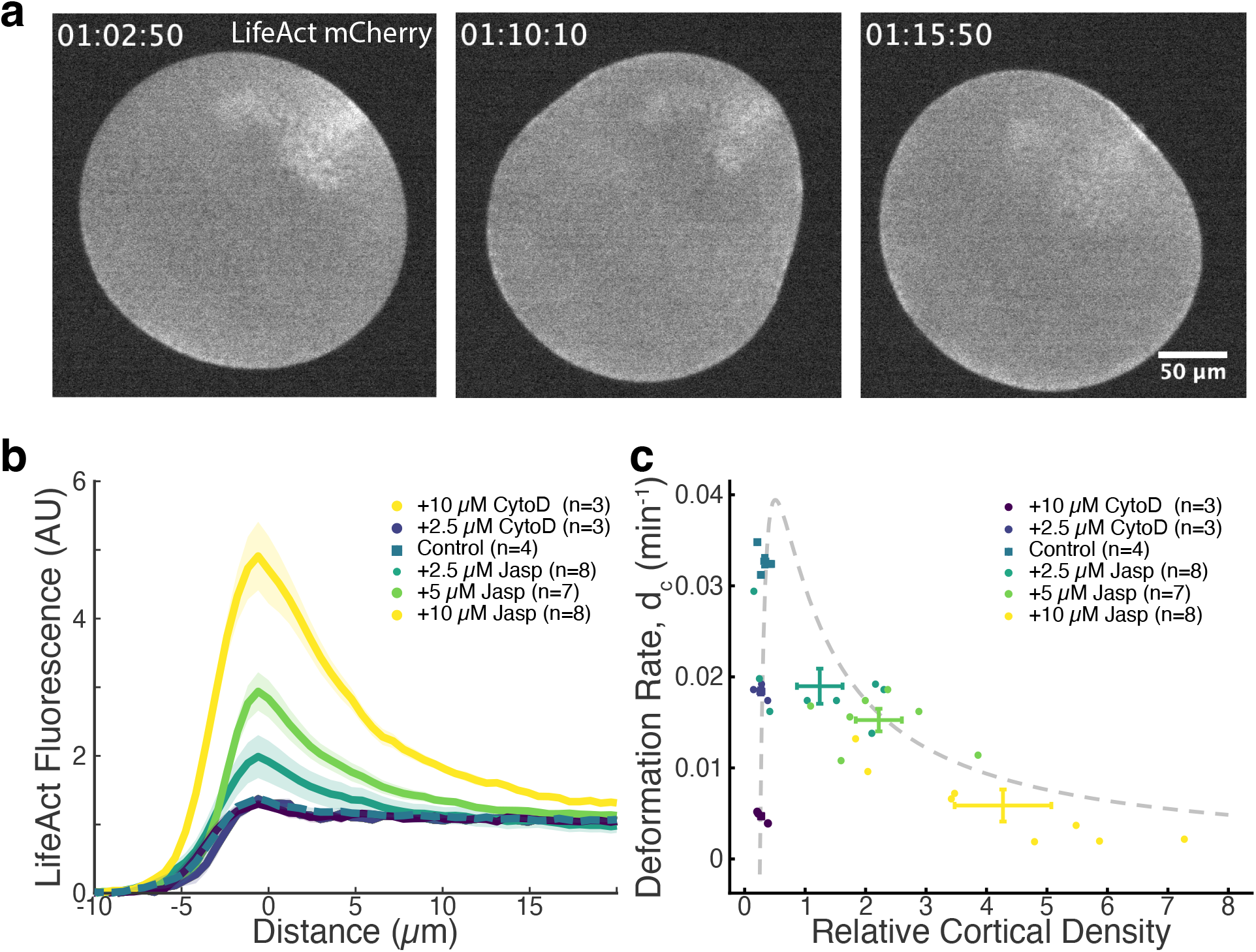
Characteristic deformation rate is maximized at intermediate actin density (a) LifeAct-mCherry imaging of F-actin localization during maturation (b) Average normalized radial line profiles of LifeAct-mCherry fluorescence for varying cytochalasin D and jasplakinolide concentrations (mean ± s.e.m.). n: number of oocytes per treatment condition. For each oocyte, two fluorescence linescans are taken and included in the average (c) Characteristic deformation rate, *d*_*c*_, as a function of relative cortical density. Errorbars: mean ± s.e.m. for each treatment condition. Grey dashed line: model fit. n: number of oocytes per treatment condition. Data was collected over 7 days using oocytes from 7 clutches.

To characterize the cortical actin density, line scans of LifeAct fluorescence were measured midway between the animal and vegetal poles when the SCW passed through this region, and the intensity profiles interior to the cell were fit to a decaying exponential function *I*(*r*) = *I*_0_*e*^−*r/λ*^ + *I*_*C*_, where *r* is the distance from the cell’s edge. The relative cortical density (RCD) was then calculated from these fitting parameters as *RCD* = *I*_0_*/*(*I*_*C*_ − *I*_*BG*_), where *I*_*BG*_ is the average fluorescence signal exterior to the cell (see *Materials and Methods*). A similar exponential decay was seen when Utrophin-mEGFP was used to visualize the cortical actin network (Extended Data Figure 2). Measurements of the relative cortical density and characteristic radial deformation rate were performed for individual oocytes treated with varying concentrations of cytochalasin D or jasplakinolide, allowing a direct comparison between the effects of these two treatments (Fig. 2b, Extended Data Figure 3). While the RCD for cytochalasin treated oocytes is decreased relative to wild type conditions (RCD = 0.27 ± 0.07 and 0.27 ± 0.06 for 2.5 and 10 *µ*M cytochalasin D respectively, compared to RCD = 0.33 ± 0.06 for WT, mean ± s.e.m.), differences in the RCD between the two cytochalasin conditions cannot be resolved, presumably due to the low absolute RCD for these conditions. As anticipated from the measured dose response curves (Fig. 1e,f), we find that the characteristic deformation rate is not a monotonic function of cortical actin density, but instead sharply increases before slowly decreasing with increasing cortical actin density, with a peak near the wild-type density (Fig. 2c). We note that the characteristic deformation rate for control cells, 0.031±0.001 min^−1^ (mean ± s.e.m, n=4 oocytes), is approximately two fold larger than we saw previously. We suspect this difference arises from environmental differences between the microscopy facilities used for these experiments.

Previous work has shown that on short timescales, treatment with agents targeting actin polymerization can feedback on the cortical localization of active Rho [30]. As the SCW in this system is driven by Rho activity, if treatment with either cytochalasin D or jasplakinolide decreases the enrichment of active Rho during the wave, then decreased Rho enrichment could explain the decrease in deformation rate seen for these treatments. To test this hypothesis, we overexpressed rGBD-GFP in oocytes as a proxy for active Rho, and treated oocytes with either 10*µ*M cytochalasin D or 10*µ*M jasplakinolide before maturation. As expected, rGBD-GFP was found to enrich in the cortex. To quantify the enrichment of rGBD-GFP, line scans of fluorescence intensity were collected and analyzed similar to LifeAct fluorescence, and a Relative Cortical Enrichment was calculated. rGBP-GFP was found to enrich to a similar degree for control and cytochalasin d treated oocytes and to a significantly greater degree for jasplakinolide treated oocytes (Extended Data Figure 4). Thus, the decrease in contraction rate seen for these treatments, and hence the peak in contraction rate as a function of relative cortical density cannot be attributed to a decrease in the enrichment of active Rho.

### Modeling the cell cortex as a thin active viscous shell links deformation rates to material properties

To understand the origin of the observed dependence of the radial deformation rate on actin density, we first asked how the physical properties of the cell cortex changed throughout the perturbation experiments. The force balance of the cortical material in bulk reads,

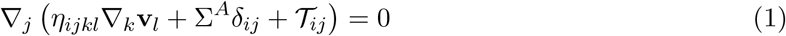

where *η*_*ijkl*_ is the viscosity tensor where the nonzero elements take the form *η*_*ijkl*_ = *ηξ*_*ijkl*_, where *η* is the magnitude of the dominant component of the viscosity and *ξ*_*ijkl*_ encodes geometric information and scalings, Σ^*A*^ is the active stress generated by interactions between actin filaments, molecular motors, and passive crosslinkers, and 𝒯 is a passive stress arising from surface tension. Einstein’s convention of summation over repeated indices is implied.

To identify how the material properties of the cell cortex change, we developed a theory for a deformable cell by mapping Eqn. 1 onto a thin shell of material (see Supplementary Information), similarly to[45–48], and used the equations of motion that we thus obtained to infer the ratio between active stresses and viscosities from experimental data. Our procedure starts by first extracting the shape *R*(*θ, t*) and its normal (*V*_*z*_) and tangential (*V*_*s*_) surface velocities from each movie (Fig. 3a, see Supplemental sections 1 and 2 for details). These quantities are then inserted into the thinshell equations of motion to determine the amplitude ||Σ^*A*^*/η*||. The results are shown in Fig. 3b (top panels), and demonstrate that the maximum value of the stress-viscosity ratio is highest for wildtype oocytes and decreases for both cytochalasin and jasplakinolide treatments. This mirrors the behavior of the characteristic deformation rate *d*_*c*_ itself (Fig. 2c).

**Figure 3:**
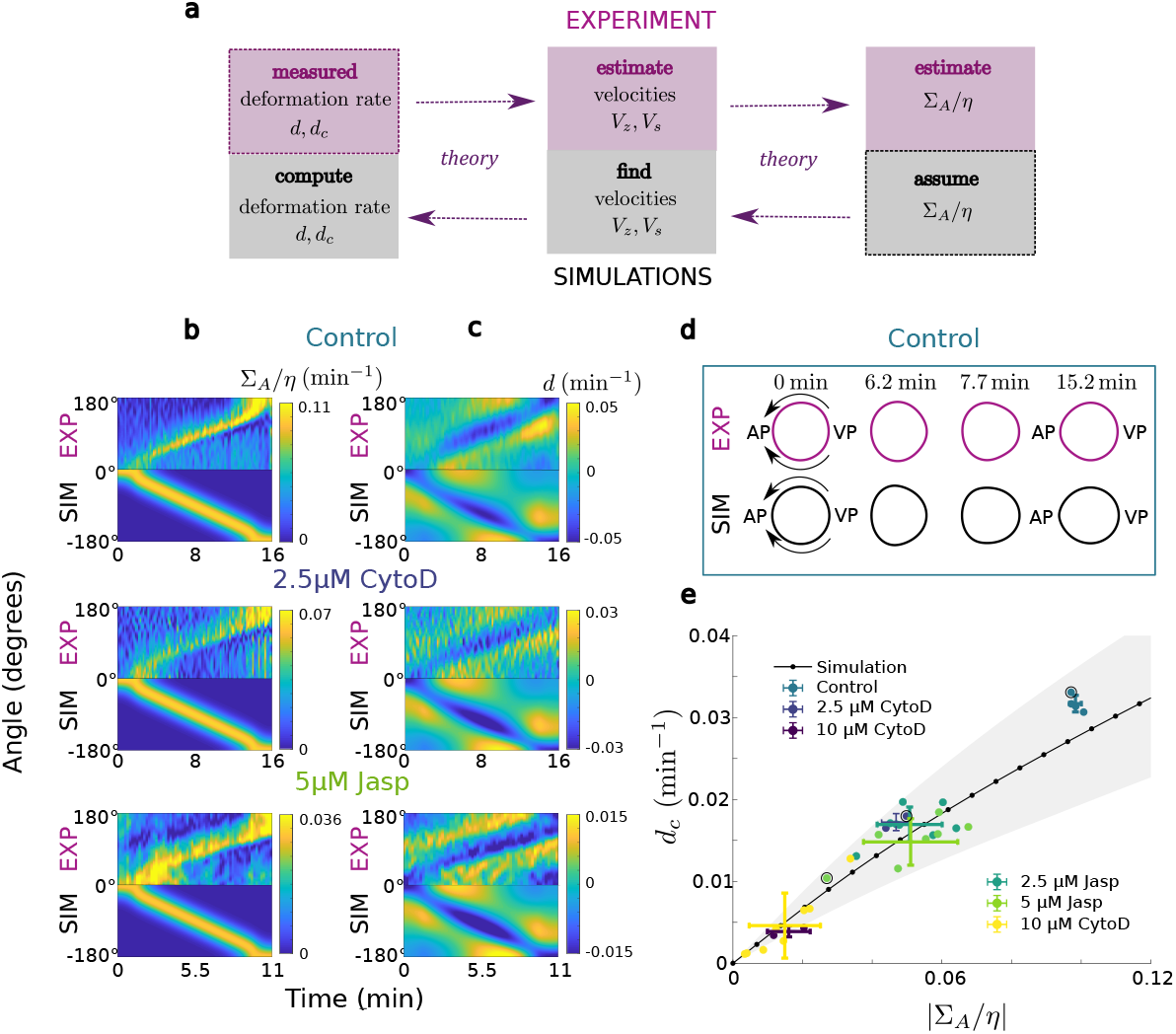
Using the active surface model to quantitatively understand experiments. (a) Description of steps followed to extract the active stress/viscosity ratio Σ_*A*_*/η* and the deformation rate *d* from experiments and simulations respectively, within our active surface model theory, discussed in detail in the SI. (b) Comparison of the estimated experimental and assumed simulation active stress Σ_*A*_*/η* kymograph (c) Comparison of the measured experimental and the computed simulation deformation rate *d* kymograph for different cases. The corresponding active stresses are shown in column (b). (d) Exemplary experimental and simulation oocyte shapes for different points in time corresponding to the control case. (e) Plot of the characteristic deformation rate *d*_*c*_ against the characteristic amplitude of active stress |Σ_*A*_*/η*| obtained in a similar way as *d*_*c*_ is derived from *d*. Shown are both the simulation values, as well as data points of representative experiments. The vertical errors for simulations (gray area) stand for the standard deviation of the minimum values of *d* in time. The vertical and horizontal errors for the experimental data points stand for the standard deviation in *d*_*c*_ and |Σ_*A*_*/η*| for different datasets for each color-coded drug treatment. The experimental points surrounded by the black circle are the ones used for comparison of simulations and experiment in columns (b) and (c). The parameter values used for the simulation are shown in Table 1 of the Supplementary Information.

To further test this behavior, we next solved for the shape changes expected in a cell with a given stress-viscosity ratio as a function of time. While the surface contraction wave has been shown to be driven by an active band of Rho [31], we here focus on the mechanics of deformation and implicitly include the signaling dynamics of Rho itself by choosing the width of the active Rho band and its traveling speed to take values consistent with experimental measurements (see Supplementary Information Table 1). In Fig. 3c (bottom panels) we report the deformation rates from these numerical experiments. We conclude that generically, the deformation rate varies almost linearly with the stress-viscosity ratio, as expected [45] (Fig. 3e) and thus that our pharmacological treatments altered the stress-viscosity ratio of the cell cortex. We next sought to understand the filament level origins of this cortical scale effect.

### An active fluid model coarse-grained from microscopic interactions

To obtain predictions for the dependence of *η* and Σ_*A*_ on the actin density, we adapted a recently developed theoretical framework which allows a microscopic description of the system to be coarsegrained into an emergent mechanical model [32]. An application of this general framework to the actomyosin system considered here begins with a simplified microscopic description of the system and considers three elements: actin filaments, passive crosslinkers, and molecular motors.

Molecular motors and passive crosslinkers exert forces between the filaments which they connect. For passive crosslinkers, these forces are taken to be proportional to the velocity difference between the points on the actin filament connected by the crosslinker. Thus, for a single crosslinker bound between position *s*_*i*_ on the *i*-th actin filament and *s*_*j*_ on the *j*-th filament, the force exerted between filaments is given by

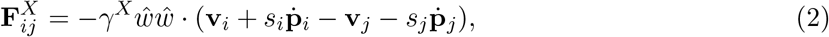

where **v**_*i*_ and **p**_*i*_ are the velocity and direction of filament *i*, and *γ*^*X*^ describes the coupling strength (see Fig. 4a). The unit vector 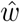 points from the between point of the crosslinks on filament *i* and *j* and is defined as 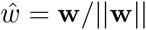 with **w** = **x**_*i*_ + *s*_*i*_**p**_*i*_ − **x**_*j*_ − *s*_*j*_**p**_*j*_. When multiple passive crosslinkers are bound between filament pairs, the net force exerted between filaments becomes,

**Figure 4:**
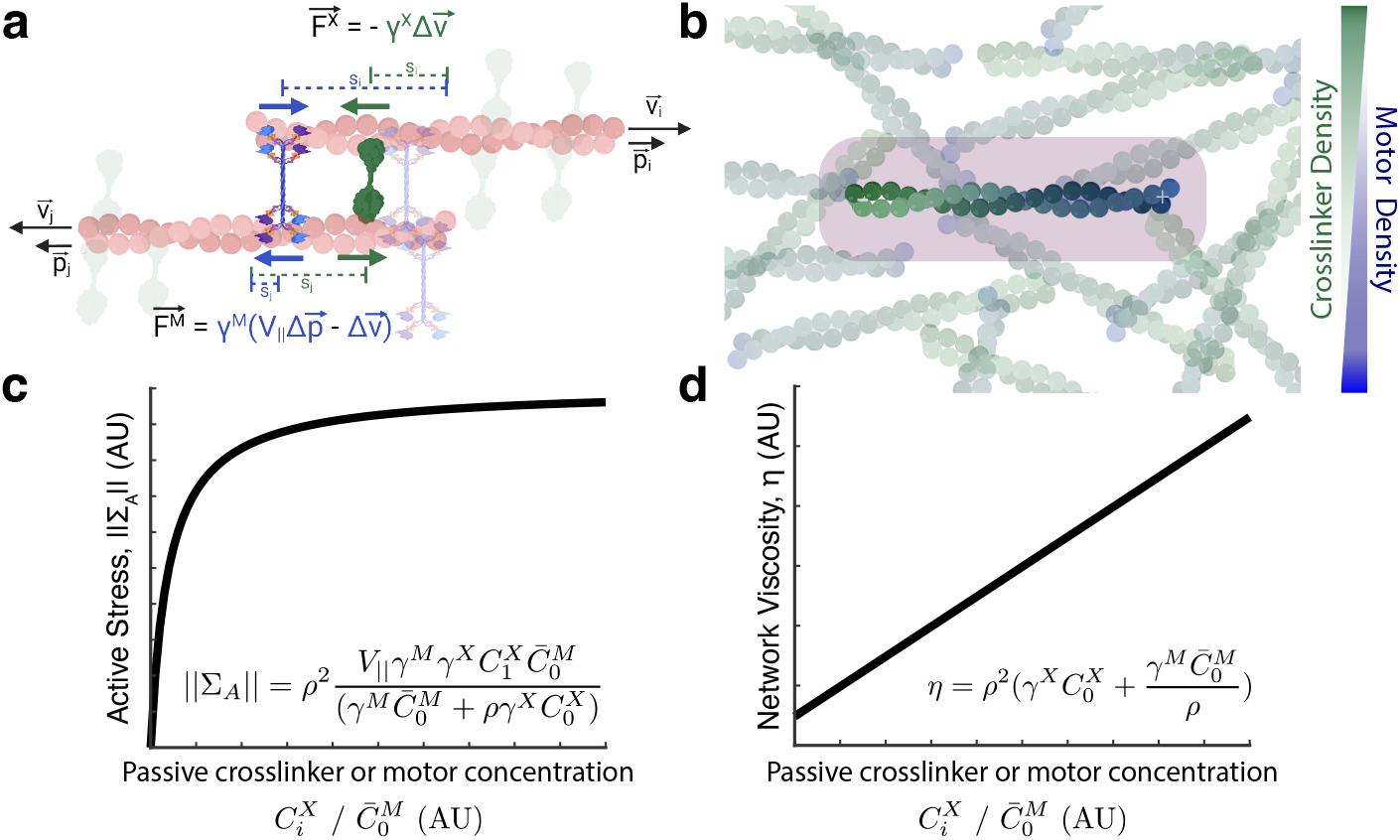
Active fluid model (a) Schematic of filament-scale forces between antiparallel actin fila-ments from individual motors (blue) and passive crosslinkers (green). (b) In the model considered in the main text, motors (blue) are uniformly distributed on actin filaments, while passive crosslinkers (green) accumulate near the filament end they motor walks away from. (c,d) Functional forms of the scalings of the active stress magnitude, ||Σ_*A*_||, (c) and the viscosity, *η*, (d) with the concentrations of either passive crosslinkers 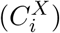 or motors 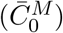.

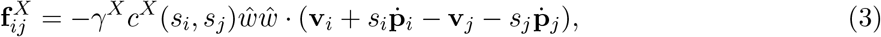

where *c*^*X*^(*s*_*i*_, *s*_*j*_) is the number of passive crosslinkers that are bound between positions *s*_*i*_ and *s*_*j*_, which can spatially vary along filaments. Motor molecules likewise contribute to the frictional coupling between filaments, but due to their stepping motion along filaments, exert additional active forces. The net force exerted by motors between filaments, 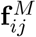 can be written as,

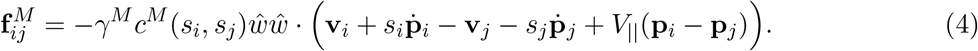

where the coefficient *V*_||_ is the unloaded speed of the motor, *γ*^*M*^ is the motor friction, and *c*^*M*^ (*s*_*i*_, *s*_*j*_) is the density of motor molecules connecting two specific filament positions. We further postulate functional forms for the motor and crosslinker densities,

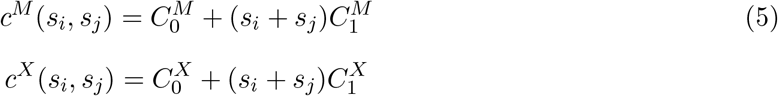

where 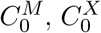 represent the number of uniformly bound motors and crosslinkers, and 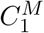 and 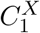 capture nonuniformity of binding along filaments. Given Eqns. 3, 4, and 5, predictions for *η* and Σ_*A*_, can be derived by integrating over all possible configurations of motors and crosslinkers [32] (Fig. 4b, see Supplementary Information). The viscosity of the system is predicted to be

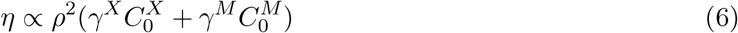

Furthermore, the active stress is predicted to be,

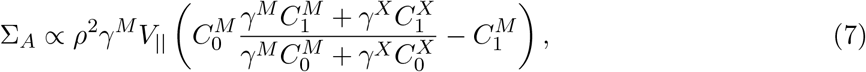

With this, the contraction rate is expected to be,

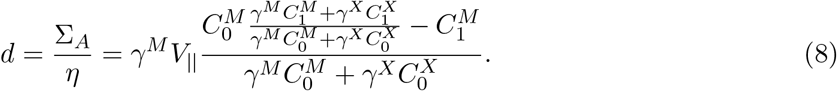

To relate these results to our experimental findings, we next need to specify how *C*^*M*^ and *C*^*X*^ change as a function of actin density, *ρ*. The average number of passive crosslinks and motor molecules bound between two filaments can depend on the overall actin density. We illustrate this by discussing two crosslinks A and B with different behaviors. Assume that the total cortical concentration of A is independent of actin filament numbers and low enough such that the binding rate of A is limited by the total concentration of A itself. As the number of filaments in the cortex is increased we expect the number of A per filament pair to drop roughly inversely proportional to the number of actin filaments. In contrast, if crosslink B is so abundant that its binding is limited by the number of available binding sites we expect a different behavior, where the number of B bound depends on the total number of binding sites available per filament, which is independent of the total number of filaments. While the binding of myosin and passive crosslinks in the cortex is likely much more complex, the key point that the average number of each type of crosslink per filament pair depends on actin density differently holds irrespectively. Importantly, since the concentration of motors and passive crosslinks per filament pair are functions of the total actin amount, *d* is a function of *ρ* as well. To capture this, we take 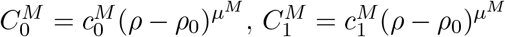, and 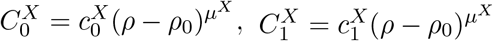. With this equation 8 becomes

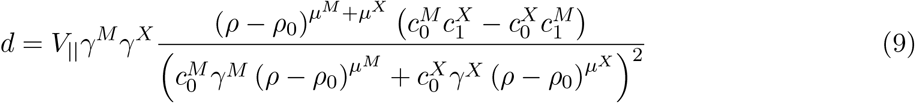

Taking |*µ*^*X*^ − *µ*^*M*^ | = 1, for simplicity, we obtain,

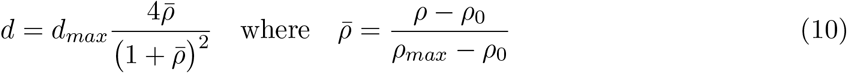

With

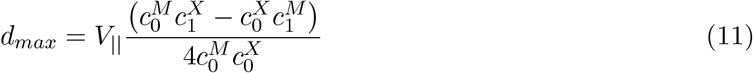

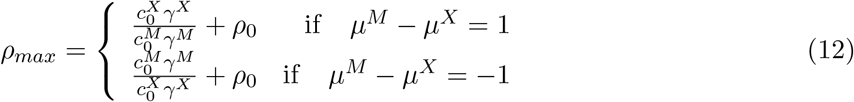

Importantly, the form of Eq. 10 is independent of microscopic assumptions. It is parameterized by the three fit parameters *d*_*max*_, *ρ*_*max*_ and *ρ*_0_. The first fit parameter *d*_*max*_ has units of contraction rate (min^−1^). It determines the height of the peak of the the density contraction curve. The second, *ρ*_*max*_ has dimensions of relative density and sets the position of the maximum. Finally, *ρ*_0_ sets the zero intercept of the curve, and could be interpreted as the percolation threshold below which the network is too sparse to be fully connected by motors and crosslinks, and thus can not contract. The sign of *d*_*max*_ determines whether the model cortex is contractile (*d*_*max*_ > 0) or extensile (*d*_*max*_ < 0). In our experiments *d*_*c*_ > 0, and thus only microscopic models with *d*_*max*_ > 0 are consistent with the data. Thus, our data could be explained by a microscopic model in which either (i) a passive crosslinkers accumulate near the end of the filament which motors walks towards 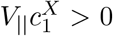, or (ii) the motor accumulates near the filament end that it walks away from 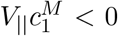; or (iii) a combination of (i) and (ii). For example, scenario (i) would be consistent with a crosslinker that tracks growing filament ends, for example a formin, some of which have been show *in vitro* to be capable of crosslinking actin filaments [49]. Scenario (ii) would be consistent with actin filament plus ends growing at a faster speed than motors walk, leading to an effective depletion of motors at the fast growing end. This effect been previously recognized to promote network contraction in an *in vitro* microtubule/motor system [50]. In any case, we stress that these two scenarios lead to equivalent predictions for the characteristic deformation rate considered here.

We next asked for which parameter combinations Eq. 10 was quantitatively consistent with the observed deformation rate changes. Fitting Eqn. 10 to the average characteristic radial deformation rate and relative cortical actin density for each treatment condition provides excellent quantitative agreement with the measured data (Fig. 2e). The underlying model parameters that best describe the data are *d*_*max*_ = 0.04 ± 0.002*µ*m min^−1^, *ρ*_*max*_ = 0.51 ± 0.02, and *ρ*_0_ = 0.25 ± 0.02 (fit values ± standard deviation). Note that models with |*µ*^*M*^ − *µ*^*X*^ | ≠ 1, failed to produce convincing fits.

### Testing and parameterizing the active fluid model through protein inhibition and overexpression experiments

In the active fluid model, forces are generated by the activity of molecular motors. To test this assumption and confirm that myosin activity underlies deformation during the SCW, oocytes were treated with the myosin inhibitor blebbistatin, which has previously been shown to almost completely suppress contraction during the SCW [31]. We find that treatment with 200 *µ*M blebbistatin substantially decreases the characteristic deformation rate, consistent with myosin’s role in driving contraction, while treatment with an inactive blebbistatin enantiomer did not significantly change the deformation rate (Extended Data Figure 5).

To further test the active fluid model, we performed experiments increasing the concentration of either a passive actin crosslinking protein or an active motor. In the active fluid model, the viscosity, *η*, and the active stress magnitude, ||Σ_*A*_|| are predicted to scale differently when the concentrations of either passive crosslinker or active motors are varied (Fig. 4c,d, Eqns. 6, 7). The concentrations of passive crosslinkers and motor proteins enter the characteristic deformation rate by modulating *ρ*_*max*_, i.e. the actin density at which contraction rates are maximized (see Eq 12).

Upon modulating the amount of passive crosslinker and motor, we expect *ρ*_*max*_ to depend on the fractional changes *f* ^*X*^, *f* ^*M*^ of the passive crosslinker and motor concentrations, respectively, as

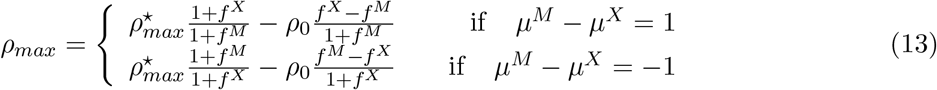

where 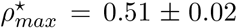 denotes the value of *ρ*_*max*_ for the case of unperturbed concentrations. Further, the density *ρ*_0_, which is related to the crosslink density at which the actin network becomes fully percolated [15], and is independent of the densities of crosslinks and motors. In what follows, we will present our fits assuming that *µ*^*M*^ − *µ*^*X*^ = −1, consistent for example with a scenario where motor binding is filament limited while crosslinker binding is crosslinker limited. We do, however, note that the second scenario provides equally good fits, and our data do not a priori allow distinguishing between the two scenarios; see SI.

In unperturbed oocytes the cortical density is close to 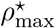 (see Fig. 2c). Thus, according to our theory, they operate near the maximum contraction rate which the system can achieve. This implies a testable prediction: the contraction rates should go down as passive crosslinkers are overexpressed and, more surprisingly, that contractions should also decrease when the motor myosin is overexpressed (see Eqn. 10,13).

To test these predictions, we first overexpressed an actin crosslinking protein in untreated oocytes. While a variety of passive actin crosslinkers localize to the cortex [51], we here use *α*-actinin as a model passive crosslinker. Fluorescent mEGFP-labeled *Patiria miniata α*-actinin [43] was over-expressed by injecting oocytes with the corresponding mRNA (see *Materials and Methods*), and we make use of the natural variability in protein expression level to assess changes in the characteristic radial deformation rate over a range of *α*-actinin concentrations.

Characteristic deformation rates and the fluorescence signals of *α*-actinin-mEGFP, which we use as a proxy for *α*-actinin concentration, were measured for individual oocytes (Fig. 5a, Supplementary Video 3). Overall, we find a general trend of decreasing deformation rate with increasing levels of *α*-actinin overexpression, qualitatively consistent with the prediction of the active fluid model. To quantitatively test the model, we take the actin density to be the previously measured wild-type value, *ρ* = *ρ*_*W T*_ and relate the measured *α*-actinin-mEGFP fluorescence signal to the fractional change in passive crosslinker concentration as, *f* ^*X*^ = *α*_*X*_*I*_*X*_. Using the values for 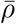, *ρ*_0_ and *d*_*max*_ determined in (Fig. 2c), we fit to the experimental data, and found quantitative agreement between theory and experiment, providing a measure of the sole fit parameter, *α*_*X*_ = 11.9 ± 5.1 (fit value ± 95% confidence interval). Thus, our active fluid model is consistent with the results of the alpha-actinin overexpression experiments.

**Figure 5:**
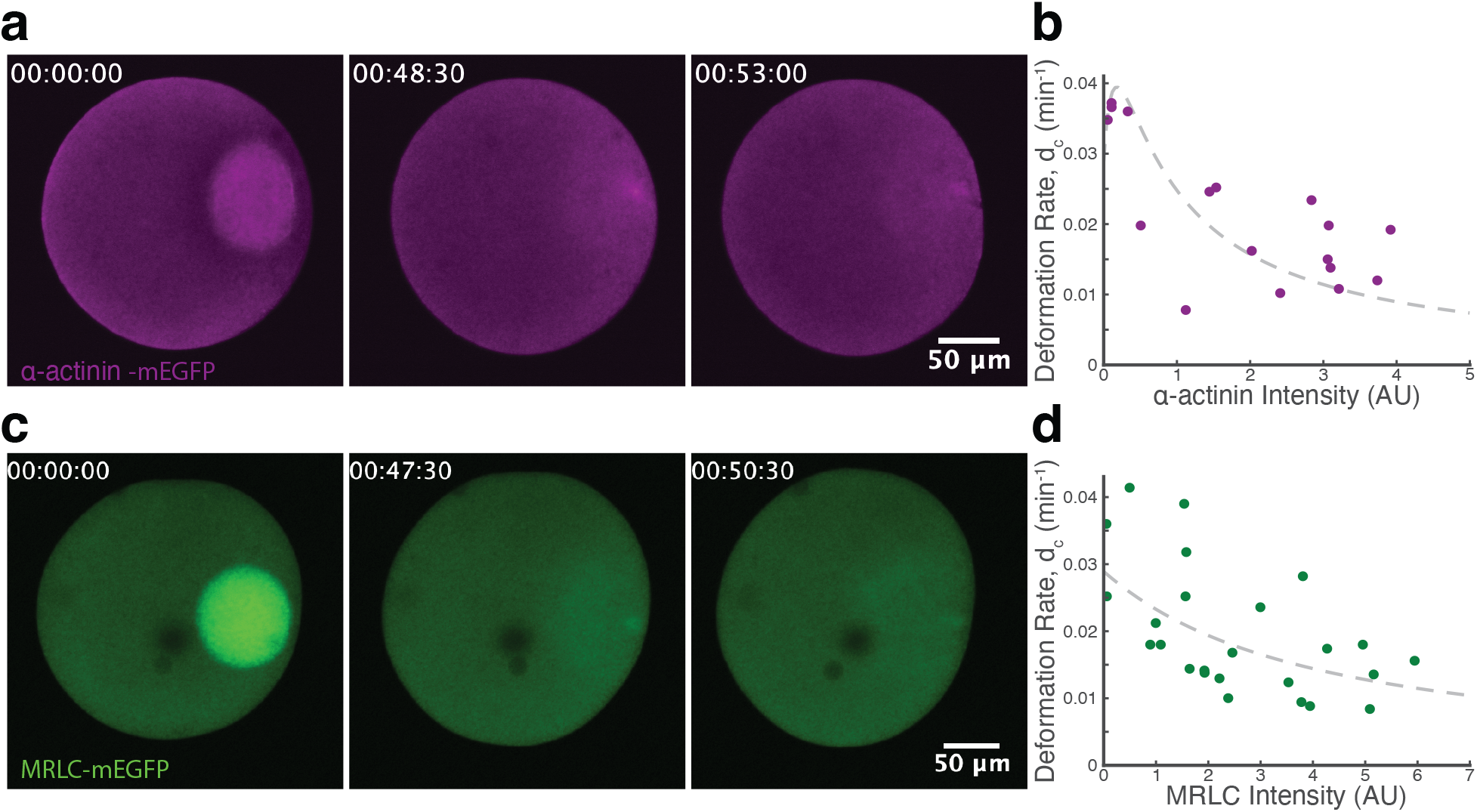
Crosslinker and motor overexpression to test active fluid model predictions. (a) Timecourse of SCW for an oocyte overexpressing *α*-actinin-mEGFP. (b) Characteristic deformation rate as a function of *α*-actinin-mEGFP intensity (n=17 oocytes). Data was collected over 3 days using oocytes from 3 clutches. Grey dashed line: model fit (c) Timecourse of SCW for an oocyte overexpressing MRLC-mEGFP. (d) Characteristic deformation rate as a function of MRLC-mEGFP intensity (n=25 oocytes). Data was collected over 4 days using oocytes from 4 clutches. Grey dashed line: model fit

The active fluid model further predicts how the characteristic deformation rate should change with the concentration of active motors. MRLC overexpression in *Patiria miniata* oocytes has previously been shown to increase the strength of the SCW [28, 31] and to increase nonequilibrium activity in the cortex [52]. In the active fluid model presented here, motors contribute both active forces and an effective friction between sliding filaments. As such, changes in the concentration of motor proteins are predicted to change both the emergent active stress and network viscosity. As motor concentration increases, the model predicts that network viscosity will grow faster than active stress, and hence the rate of deformation will decrease with increasing motor concentration (Fig. 5c,d).

To experimentally test this prediction, mEGFP-labeled *Patiria miniata* myosin regulatory light chain (MRLC) was overexpressed in oocytes (Fig. 5c, Supplementary Video 4). The characteristic deformation rate and MRLC-mEGFP fluorescence were measured for individual oocytes (see *Materials and Methods*) and the resulting data were fit to Eqns. 10, 13, under the assumption that the measured MRLC-mEGFP fluorescence signal is proportional to the fractional change in active motor concentration, *f* ^*M*^ = *α*_*M*_ *I*_*M*_ . Once again, using only a single free fitting parameter, *α*_*M*_, we find quantitative agreement between the measured data and the prediction of the active fluid model (Fig. 5d), providing a measure of *α*_*M*_ = 0.45 ± 0.25 (fit value ± 95% confidence interval). We conclude, that our model is consistent with the observed effects of Myosin overexpression. To demonstrate this, we plot all of our data in terms of 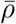, and demonstrate that two types of overexpression experiments, as well as our pharmacological perturbations, and the unperturbed data, fall onto a single master curve predicted by Eq 10 (Fig. 6)

**Figure 6:**
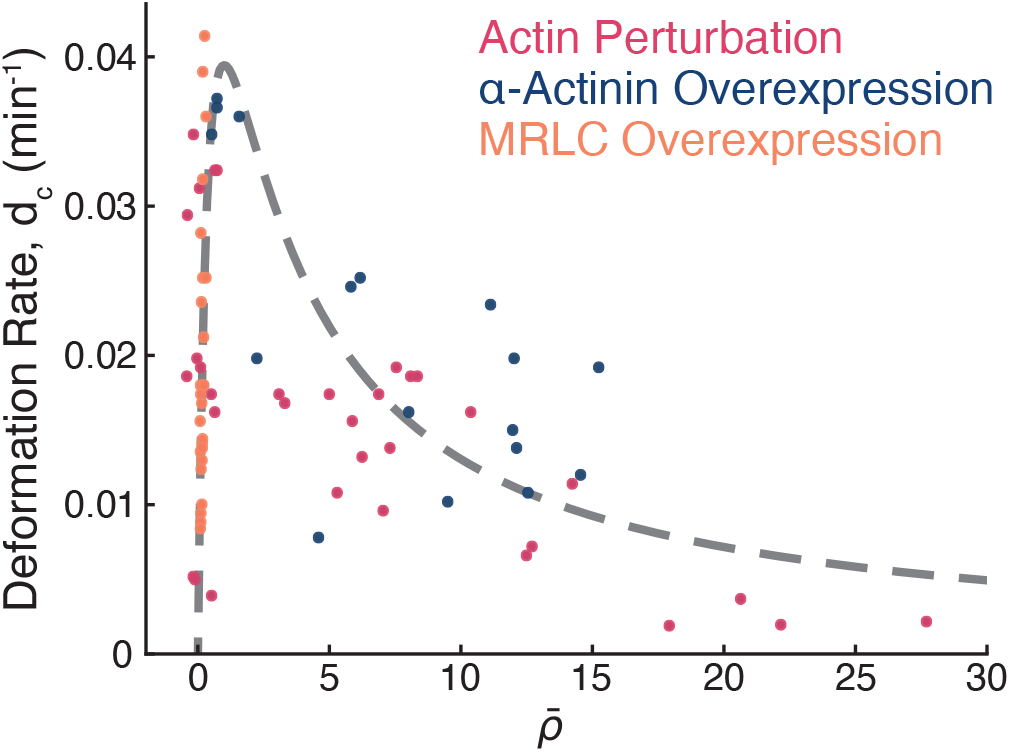
Using the fit parameters data from experiments with varying Relative Cortical Density (Fig. 2c), *α*-actinin-mEGFP concentration (Fig. 5b), and MRLC-mEGFP concentration (Fig. 5d) can be reparameterized and plotted as a function of 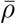. Upon replotting, the data collapse to a curve predicted by the active fluid model.

## Discussion

Here, we used surface contraction waves in maturing sea star oocytes as a model to study actomyosin contractility *in vivo*. By controlling cortical actin density, we find that the deformation rate is maximum near the wild-type density and decreases when the cortical actin density is either increased or decreased. To understand this phenomenon, we first analyzed the deformations in terms of a deformable active surface model for the oocyte, and found the observed deformation rates indicate that the ratio of cortical contractile stress to cortical viscosity was actively modulated by actin density. To better understand how this could be happening, we developed an active fluid model coarse-grained from a microscopic description of the system. The model predicts the dependence of the radial deformation rate on the relative abundance of passive crosslinkers and motors per filament pair, and shows that contraction rates are maximized for intermediate amounts of passive crosslinks. As the abundance of passive and motorized crosslinks per filament pair are functions of the actin density itself, we further predict that this implies that the maximal contraction rate is reached at intermediate cortical densities following the functional form given by Eqn. 10, which is in good quantitative agreement with our data. The theory which we obtain has only three free parameters, which are the maximum contraction rate *d*_*max*_, the actin density at which this rate is reached *ρ*_*max*_ and the minimal density *ρ*_0_ below which the cortex can not contract - presumably because it is no longer fully percolated. Strikingly *d*_*max*_ is fully set by the asymmetry of the motor and crosslink binding, and motor speeds, but is independent of absolute motor concentrations. In contrast, *ρ*_*max*_ depends crucially on concentrations, but is independent of detailed motor properties. This highlights the role that actin density itself plays for controlling cortical contractility. We thus suggest that the feedback loop that we uncovered here, allows controlling contractile instabilities, without the need for additional biochemical feedback loops. By using model parameters measured from fitting the active fluid model to the actin density, *α*-actinin overexpression, and MRLC overexpression data, data from all three sets of experiments could be reparameterized and plotted as a function of the same parameter, leading to data collapse to a curve predicted by the active fluid model.

Having established quantitative agreement between the active fluid model and the data, we now discuss the microscopic picture that underlies this model. The key microscopic ingredient is a crosslinker that clusters near filament ends and a uniformly distributed motor, whose per filament concentration scales inversely with actin density, for simplicity. We stress that other models are also consistent with the data presented here. These include models where passive crosslinkers are uniformly distributed on filaments while myosin accumulates towards the end it walks away from, and a combination of these models where myosin and passive crosslinkers accumulate at opposite ends. Our key advance is to identify the unifying features of all allowed microscopic models, thus identifying the control knobs available to cells during development. The key features are (i) microscopically broken symmetry on the filament scale, which requires either an asymmetry in the driving force (due to the spatial localization of motors) or in the frictional coupling (due to the spatial localization of passive crosslinkers) and (ii) the balance between per filament motoring and friction. Importantly, these two key aspects of the microscale physics can be independently regulated, as is demonstrated by their very different dependence on actin density. Beyond actin density, crosslinker turnover has also been shown to be a mechanism for regulating active stresses [53]. We speculate that it acts via the same two control knobs.

Similar to previous *in vitro* observations of contractile actomyosin [12, 13, 54], both the *in vivo* measurements and the active fluid model presented here show a decrease in network contractility with increasing crosslinker concentration. In contrast to previous work, the results here show that at high motor concentration, network contractility decreases, a qualitatively different behavior from both previous *in vitro* measurements [12, 55] and theoretical predictions from a filament buckling model [56], where the network contraction rate instead saturates with increasing motor concentration rather than decreasing. Network connectivity has been used to explain previous *in vitro* observations: at low connectivity motor forces cannot propagate to larger scales [15], while at high connectivity network contractility decreases due to either a substantial increase in network rigidity [1, 57] or to a decrease in filament buckling [13, 56]. The active fluid model that we developed, elucidates an alternative mechanism that can lead to this phenomenon. At low crosslinker concentrations, active stress falls off faster that viscosity, and thus the network deformation rate decreases. Experimentally measuring the network connectivity of *in vivo* actin cortices is a challenge, and in the future electron microscopy could potentially be used to measure whether our experimental results at low actin density are above the percolation threshold. Such studies could also uncover how actin perturbations in this system modulate cortical actin network architecture, which can influence contractility separately from actin density alone [8, 9, 13]. At high crosslinker concentration, the model predicts viscosity increases faster than active stress, and hence the deformation rate decreases. This idea shares similarities with models where increased network connectivity leads to high stiffness: both would emerge from a high degree of filament crosslinking. An intriguing possibility is that both classes of models are limiting cases of a universal mechanical framework. Exploring this possibility will be an exciting avenue for future research.

Finally, we note that in the system considered here, the deformation rate is maximal near the wild type composition, and perturbing the system through changing the cortical actin density, *α*-actinin concentration, or MRLC concentration largely only decreases the deformation rate. For the model considered in the main text, the maximal deformation rate is given by 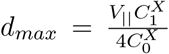, implying that increasing the deformation rate further would only be possible through either increasing the relative asymmetry of crosslinker localization or increasing the motor walking speed. Increasing the motor walking speed or changing the system composition would have energetic consequences and energetic considerations can impose additional constraints in living nonequilibrium systems [58]. Understanding the energetic constrains of the emergent dynamics could further constrain possible microscopic models, and would require going beyond network architecture towards a thermodynamic description of such living active systems [5].

## Data and Code Availability

All data that support the plots within this paper and other findings of this study are available from the corresponding authors upon reasonable request. Images were analyzed using custom written MATLAB code available at https://github.com/foster61012/Starfish_SCW.

## Author Contributions

P.J.F. and N.F. initiated the project and designed the experiments. P.J.F. and J. L. performed the experiments and analysed the experimental data. S.F. and A.Z. designed the active fluid model. All authors participated in interpreting the experimental and theoretical results and in writing the manuscript.

## Acknowledgments

This research was supported by the National Science Foundation CAREER Award to N.F.. P.J.F. acknowledges support from the Gordon and Betty Moore Foundation as a Physics of Living Systems Fellow through grant no. GBMF4513. P.J.F. acknowledges support from the NSF MRSEC DMR-2011846. S.F. and A.Z. have been funded by the Vienna Science and Technology Fund (WWTF) through grant no. [10.47379/VRG20002]. Fig. 4a,b and Fig. S6 were created with BioRender.com.

## References

1. Banerjee, S., Gardel, M. L. & Schwarz, U. S. The Actin Cytoskeleton as an Active Adaptive Material. Annu Rev Condens Matter Phys 11, 421–439 (2020).

2. Foster, P. J., Fürthauer, S., Shelley, M. J. & Needleman, D. J. From cytoskeletal assemblies to living materials. Curr. Opin. Cell Biol. 56, 109–114 (2019).

3. Marchetti, M. C. et al. Hydrodynamics of soft active matter. Reviews Of Modern Physics 85, 1143–1189 (2013).

4. Needleman, D. & Dogic, Z. Active matter at the interface between materials science and cell biology. Nature reviews materials 2, 1–14 (2017).

5. Bowick, M. J., Fakhri, N., Marchetti, M. C. & Ramaswamy, S. Symmetry, Thermodynamics, and Topology in Active Matter. Phys. Rev. X 12, 010501 (Feb. 2022).

6. Lenz, M. Reversal of contractility as a signature of self-organization in cytoskeletal bundles. eLife 9, e51751 (2020).

7. Wollrab, V. et al. Polarity sorting drives remodeling of actin-myosin networks. Journal of Cell Science 132, jcs219717–14 (2018).

8. Koenderink, G. H. & Paluch, E. K. Architecture shapes contractility in actomyosin networks. Current Opinion in Cell Biology 50, 79–85 (2018).

9. Chugh, P. et al. Actin cortex architecture regulates cell surface tension. Nature Cell Biology 19, 689–697 (2017).

10. Feld, L. et al. Cellular contractile forces are nonmechanosensitive. Science Advances 6, eaaz6997 (Apr. 2020).

11. Backouche, F., Haviv, L., Groswasser, D. & Bernheim-Groswasser, A. Active gels: dynamics of patterning and self-organization. Physical biology 3, 264–273 (2006).

12. Bendix, P. M. et al. A Quantitative Analysis of Contractility in Active Cytoskeletal Protein Networks. Biophysical Journal 94, 3126–3136 (2008).

13. Ennomani, H. et al. Architecture and Connectivity Govern Actin Network Contractility. Current Biology 26, 616–626 (Mar. 2016).

14. Tan, T. H. et al. Self-organized stress patterns drive state transitions in actin cortices. Science Advances 4, eaar2847 (2018).

15. Alvarado, J., Sheinman, M., Sharma, A., MacKintosh, F. C. & Koenderink, G. H. Molecular motors robustly drive active gels to a critically connected state. Nat. Phys. 9, 591–597 (Aug. 2013).

16. Murrell, M. P. & Gardel, M. L. F-actin buckling coordinates contractility and severing in a biomimetic actomyosin cortex. Proceedings Of The National Academy Of Sciences Of The United States Of America 109, 20820–20825 (2012).

17. Köhler, S., Schmoller, K. M., Crevenna, A. H. & Bausch, A. R. Regulating contractility of the actomyosin cytoskeleton by pH. Cell Rep. 2, 433–439 (2012).

18. Murrell, M., Oakes, P. W., Lenz, M. & Gardel, M. L. Forcing cells into shape: the mechanics of actomyosin contractility. Nature Reviews Molecular Cell Biology 16, 486–498 (2015).

19. Lenz, M. Geometrical Origins of Contractility in Disordered Actomyosin Networks. Phys. Rev. X 4, 041002 (2014).

20. Kruse, K. & Jülicher, F. Actively contracting bundles of polar filaments. Phys. Rev. Lett. 85, 1778–1781 (2000).

21. Foster, P. J., Fürthauer, S., Shelley, M. J. & Needleman, D. J. Active contraction of microtubule networks. eLife 4, e10837 (2015).

22. Torisawa, T., Taniguchi, D., Ishihara, S. & Oiwa, K. Spontaneous Formation of a Globally Connected Contractile Network in a Microtubule-Motor System. Biophys. J. 111, 373–385 (2016).

23. Tan, R., Foster, P. J., Needleman, D. J. & McKenney, R. J. Cooperative Accumulation of Dynein-Dynactin at Microtubule Minus-Ends Drives Microtubule Network Reorganization. Developmental Cell 44, 233–247 (2018).

24. Lenz, M., Thoresen, T., Gardel, M. L. & Dinner, A. R. Contractile units in disordered actomyosin bundles arise from F-actin buckling. Physical Review Letters 108, 238107 (2012).

25. Chen, S., Markovich, T. & MacKintosh, F. C. Motor-Free Contractility in Active Gels. Phys. Rev. Lett. 125, 208101 (2020).

26. Chen, S., Markovich, T. & MacKintosh, F. C. Motor-free contractility of active biopolymer networks. arXiv preprint arXiv:2204.00222 (2022).

27. Lénárt, P. et al. A contractile nuclear actin network drives chromosome congression in oocytes. Nature 436, 812–818 (July 2005).

28. Bun, P., Dmitrieff, S., Belmonte, J. M., Nédélec, F. J. & Lénárt, P. A disassembly-driven mechanism explains F-actin-mediated chromosome transport in starfish oocytes. eLife 7, e31469 (2018).

29. Kučera, O. et al. Anillin propels myosin-independent constriction of actin rings. Nat. Commun. 12, 4595 (2021).

30. Bement, W. M. et al. Activator–inhibitor coupling between Rho signalling and actin assembly makes the cell cortex an excitable medium. Nature cell biology 17, 1471–1483 (2015).

31. Bischof, J. et al. A cdk1 gradient guides surface contraction waves in oocytes. Nature Communications 8, 1–10 (2017).

32. Fürthauer, S., Needleman, D. J. & Shelley, M. J. A design framework for actively crosslinked filament networks. New Journal of Physics 23, 013012 (2021).

33. Fürthauer, S. et al. Self-straining of actively crosslinked microtubule networks. Nature physics 15, 1295–1300 (2019).

34. Klughammer, N. et al. Cytoplasmic flows in starfish oocytes are fully determined by cortical contractions. PLoS Computational Biology 14, e1006588 (2018).

35. Wigbers, M. C. et al. A hierarchy of protein patterns robustly decodes cell shape information. Nature Physics 17, 578–584 (2021).

36. Hara, K. Cinematographic observation of “surface contraction waves” (SCW) during the early cleavage of axolotl eggs. Wilhelm Roux’ Archiv für Entwicklungsmechanik der Organismen 167, 183–186 (1971).

37. Satoh, N. ‘Metachronous ‘ cleavage and initiation of gastrulation in amphibian embryos. Development, Growth and Differentiation 19, 111–117 (1977).

38. Lewis, C. A. Ultrastructure of a fertilized barnacle egg (Pollicipes polymerus) with peristaltic constrictions. Wilhelm Roux’ Archiv für Entwicklungsmechanik der Organismen 181, 333–355 (1977).

39. Sardet, C., Speksnijder, J., Inoue, S. & Jaffe, L. Fertilization and ooplasmic movements in the ascidian egg. Development 105, 237–249 (1989).

40. Liu, J. et al. Light-induced cortical excitability reveals programmable shape dynamics in starfish oocytes. Nature Physics, 1–10 (2025).

41. Narumiya, S., Tanji, M. & Ishizaki, T. Rho signaling, ROCK and mDia1, in transformation, metastasis and invasion. Cancer and Metastasis Reviews 28, 65–76 (2009).

42. Bubb, M. R., Senderowicz, A. M., Sausville, E. A., Duncan, K. L. & Korn, E. D. Jasplakinolide, a cytotoxic natural product, induces actin polymerization and competitively inhibits the binding of phalloidin to F-actin. Journal of Biological Chemistry 269, 14869–14871 (1994).

43. Mori, M. et al. An Arp2/3 Nucleated F-Actin Shell Fragments Nuclear Membranes at Nuclear Envelope Breakdown in Starfish Oocytes. Current Biology 24, 1421–1428 (2014).

44. Wesolowska, N. et al. Actin assembly ruptures the nuclear envelope by prying the lamina away from nuclear pores and nuclear membranes in starfish oocytes. eLife 9, e49774 (2020).

45. Mietke, A., Jülicher, F. & Sbalzarini, I. F. Self-organized shape dynamics of active surfaces. Proc Natl Acad Sci U S A 116, 29–34 (Jan. 2019).

46. Bächer, C., Khoromskaia, D., Salbreux, G. & Gekle, S. A Three-Dimensional Numerical Model of an Active Cell Cortex in the Viscous Limit. Frontiers in Physics 9, 753230 (2021).

47. Borja da Rocha, H., Bleyer, J. & Turlier, H. A viscous active shell theory of the cell cortex. Journal of the Mechanics and Physics of Solids 164, 104876 (2022).

48. Wittwer, L. D. & Aland, S. A computational model of self-organized shape dynamics of active surfaces in fluids. Journal of Computational Physics: X 17, 100126 (2023).

49. Esue, O., Harris, E. S., Higgs, H. N. & Wirtz, D. The filamentous actin cross-linking/bundling activity of mammalian formins. Journal of molecular biology 384, 324–334 (2008).

50. Roostalu, J., Rickman, J., Thomas, C., Nédélec, F. & Surrey, T. Determinants of polar versus nematic organization in networks of dynamic microtubules and mitotic motors. Cell 175, 796–808 (2018).

51. Chugh, P. & Paluch, E. K. The actin cortex at a glance. Journal of Cell Science 131, jcs186254 (2018).

52. Tan, T. H. et al. Scale-dependent irreversibility in living matter. arXiv, arXiv:2107.05701v1 [physics.bio–ph] (2021).

53. Hiraiwa, T. & Salbreux, G. Role of turnover in active stress generation in a filament network. Physical Review Letters 116, 188101 (2016).

54. Janson, L. W., Kolega, J. & Taylor, D. L. Modulation of contraction by gelation/solation in a reconstituted motile model. J. Cell Biol. 114, 1005–1015 (1991).

55. Murrell, M. & Gardel, M. L. Actomyosin sliding is attenuated in contractile biomimetic cortices. Mol. Biol. Cell 25, 1845–1853 (2014).

56. Belmonte, J. M., Leptin, M. & Nédélec, F. A theory that predicts behaviors of disordered cytoskeletal networks. Mol. Syst. Biol. 13, 941 (2017).

57. Gardel, M. L. et al. Elastic behavior of cross-linked and bundled actin networks. Science 304, 1301–1305 (2004).

58. Yang, X. et al. Physical bioenergetics: Energy fluxes, budgets, and constraints in cells. Proc. Natl. Acad. Sci. U. S. A. 118, e2026786118 (June 2021).

