## Supplementary Information for "Active mechanics of sea star oocytes"

**1 Methods and Materials** **2**

### 1 Methods and Materials

#### 1.1 Sea star oocyte preparation

*Patiria miniata* were purchased from South Coast Bio-Marine and oocytes were collected as previously described [1]. Briefly, ovaries were collected through a small incision in the oral side of the sea star and fragmented in calcium-free seawater using scissors to release oocytes. Oocytes were washed twice with calcium-free seawater and kept in filtered seawater at 15°C until use. Oocytes were used within 2 days of oocyte extraction. For experiments with cytochalasin D (Abcam), jasplakinolide (Abcam), or blebbistatin (Sigma), drugs were first added to filtered sea water before being diluted to their final concentration with filtered sea water containing oocytes. Oocytes were incubated in this solution for at least 1 hour prior to maturation. Maturation was induced by the addition of 1-MA to a final concentration of 20 $\mu$M. Oocytes were incubated for 5 minutes before being placed into a flow cell made from 100$\mu$m 3M Polyester Double Sided Film Tape and sealed with Valap.

#### 1.2 In vitro mRNA transcription and microinjection

The plasmids for $\alpha$-actinin-mEGFP [2] and MRLC-mEGFP [3] were generous gifts from Péter Lénárt, and the plasmid for LifeAct-mCherry was a generous gift from Cynthia A. Bradham. Plasmids were isolated using a Miniprep kit (QIAGEN) before linearization using the appropriate restriction enzymes. *In vitro* transcription was performed using either an mMESSAGE mMACHINE T7 ULTRA Transcription Kit (for $\alpha$-actinin-mEGFP and MRLC-mEGFP) or an mMESSAGE mMACHINE SP6 Transcription Kit (for LifeAct-mCherry), including poly(A) tailing. Oocytes were microinjected with mRNA and incubated overnight at 15°C before use.

#### 1.3 Microscopy

Brightfield microscopy was performed using a Leica DM IL LED microscope equipped with an AmScope MU500-CK camera controlled using $\mu$Manager software [4] and a 10x, 0.25 NA objective. Fluorescence measurements were performed using a spinning disk confocal system consisting of a Zeiss AxioVert 200M inverted microscope, a Yokogawa CSU-22 spinning disk scan head, a 10x, 0.30 NA objective, and a Hamamatsu Orca-ER camera controlled using MetaMorph software. All microscopy was performed at room temperature.

#### 1.4 Data analysis

Images were analyzed using custom written MATLAB code available at, `https://github.com/foster61012/Starfish_SCW`.

##### 1.4.1 Characteristic Deformation Rate

To measure the characteristic deformation rate during the surface contraction wave, each frame in a video was first intensity thresholded to create an initial mask for the oocyte. Only for blebbistatin experiments, a Gaussian filter of width 2 pixels was applied prior to thresholding. Holes in this initial mask were filled in, the edges were eroded, the largest object in the resulting initial mask was taken as an image mask, and the edges were dilated. Only for blebbistatin experiments, a second round of edge erosion was performed prior to dilation. From this image mask, the x-y coordinates of the oocyte's outer contour were found by calculating the gradient of the image mask, and finding the x-y coordinates where this gradient was greater than zero. These coordinates were fit to a circle before being converted into polar coordinates, r and $\theta$, where r=0 corresponds to the circle's center. To smooth this outer contour, these r($\theta$) coordinates were fit to a $50^{th}$ order polynomial. This polynomial which was then evaluated for

500 evenly divided values for $\theta$ between 0 and $2\pi$. Substantial artifacts in the polynomial fitting were evident for small and large $\theta$ values, and thus values of the radial distance to the oocyte's outer contour, $R(\theta)$ were only retained for values of $\theta$ where $\frac{\pi}{2} < \theta < \frac{3\pi}{2}$. To find the remaining distances, $\pi$ was added to the $\theta$ values for nonsmoothed r($\theta$) coordinates, the polynomial fitting and evaluation procedure was repeated, and $\pi$ was subtracted from the evaluated angles, giving $R(\theta)$ for $0 < \theta < \frac{\pi}{2}$ and $\frac{3\pi}{2} < \theta < 2\pi$ in the original coordinates.
This procedure was repeated for each frame to build the distance between the oocyte's center for each angle and time, $R(\theta, t)$, which was then smoothed in time using a 5-point moving average. To calculate the local deformation rate, for each value of $\theta$ the temporal average of $R(\theta, t)$, $\langle R(\theta)\rangle_t$, was first calculated, and used to normalize $R(\theta, t)$, giving

$$\epsilon(\theta, t) = \frac{R(\theta, t)}{\langle R(\theta)\rangle_t} \tag{1}$$

From this, the local deformation rate was calculated by taking a time derivative as,

$$d(\theta, t) = \frac{\epsilon(\theta, t + \Delta t) - \epsilon(\theta, t)}{\Delta t} \tag{2}$$

where $\Delta t$ is the time interval between subsequent frames in the video. An example kymograph of $d(\theta, t)$ is shown in the main text (Fig. 1c). From $d(\theta, t)$, the characteristic deformation rate $d_c$ was calculated by taking magnitude of the minimum value of $d(\theta, t)$ across time (Fig. 1d) and averaging across all angles,

$$d_c = \langle |\min_{\{t\}} \big(d(\theta, t)\big)|\rangle_\theta \tag{3}$$

##### 1.4.2 SCW Propagation Speed

To measure the SCW propagation speed, kymographs of the local deformation rate, $\frac{d(}{d\theta}, t)$, were generated for each oocyte undergoing a SCW (Fig. 1c). From these kymographs, $\phi$, the angle of the propagating SCW in the $\theta$-t plane, was manually measured using Fiji. This angle was converted to a propagation speed using the relation,

$$v = R\tan(\phi) \tag{4}$$

where $R$ is the oocyte radius.

##### 1.4.3 Dose Response Curve Fitting

For the cytochalasin D titration (Fig. 2a), the value for half-maximal inhibitory concentration, $IC_{50}$, was found by fitting the mean characteristic deformation rate for each cytochalasin D treatment condition to the dose response curve,

$$d_c = \frac{d_{max}}{1 + \left(\frac{[\text{Cytochalasin D}]}{\text{IC}_{50}}\right)^n} \tag{5}$$

providing measurements of the fit parameters $d_{max} = 0.016 \pm 0.003$ min$^{-1}$, $IC_{50}$=$4.4 \pm 2.0\mu$M, and $n = 1.7 \pm 1.1$ (fit parameters $\pm$ 95% confidence intervals). Similarly, data for the jasplakinolide titration (Fig. 2b) were fit to the dose response curve,

$$d_c = \frac{d_{max}}{1 + \left(\frac{[\text{Jasplakinolide}]}{\text{IC}_{50}}\right)^n} \tag{6}$$

providing measurements of the fit parameters $d_{max} = 0.016 \pm 0.004$ min$^{-1}$, $IC_{50}$=$1.3 \pm 0.8\mu$M, and $n = 1.5 \pm 0.8$ (fit parameters $\pm$ 95% confidence intervals). $0\mu$M controls, where oocytes were treated with a volume of DMSO equivalent to the volume of added cytochalasin d or jasplakinolide, were shared for the two titrations and included in both fits.

##### 1.4.4 MRLC-mEGFP and $\alpha$-actinin-mEGFP Fluorescence Signal

The fluorescence signal from overexpressed fluorescently-tagged proteins was analyzed from the initial frame in a timelapse. Briefly, a binary mask for the oocyte was found by first calculating the intensity gradient of the initial frame. A threshold was applied to the image gradient, allowing the oocyte's edges to be identified. Points inside the edges were converted to 1's using MATLAB's imfill function, resulting in a mask that was 1 for points interior to the oocyte and 0 for points outside. The background fluorescence was taken as the average fluorescence signal for points outside of the oocytes. This background fluorescence was subtracted from the fluorescence intensity of all points interior to the oocyte. Finally, the background subtracted intensities of points interior to the oocyte were averaged and normalized by the camera exposure time, giving the $\alpha$-actinin and MRLC Intensities plotted in Fig. 4b, d.

##### 1.4.5 Relative Cortical Density

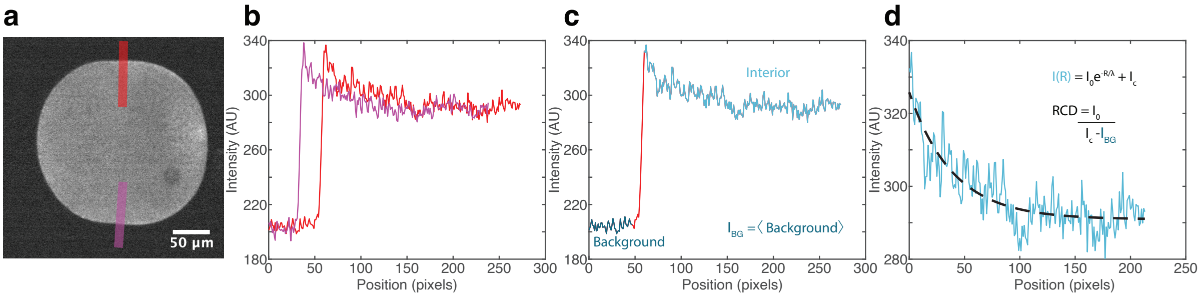

Figure S1: Relative cortical density (RCD) measurement. (a) Linescans of fluorescence intensity are taken through the cortex when the SCW is halfway between the animal and vegetal poles. (b) Example fluorescence intensity profiles. (c) Fluorescence profiles are divided into regions exterior to the oocyte (dark blue) and interior to the oocyte (light blue). The average fluorescence signal exterior to the oocyte is taken as a background intensity, $I_{BG}$ (d) The fluorescence signal interior to the oocyte is fit to a decaying exponential function, and the fit parameters are used to calculate the RCD.

To measure the relative cortical density (RCD) for oocytes expressing LifeAct-mCherry, two 20 pixel wide linescans of fluorescence intensity were taken, one on each side of the oocyte, in the radial direction from a point outside of the oocyte to a point on the interior at the time when the SCW passes midway across the oocyte (Fig. S1a,b). For each of these linescans, a background fluorescence value, $I_{BG}$ was determined by finding the point where the intensity is maximum and averaging the intensities for points outside of the oocyte and >15 pixels away from the maximum location (Fig. S1c). Intensity values interior to the oocyte starting from 2 points exterior to the maximum location were fit to an exponential function of the form,

$$I(R) = I_0 e^{-R/\lambda} + I_C \tag{7}$$

where $I_0$ is the peak cortical fluorescence signal, $\lambda$ is the length scale of the fluorescence decay, and $I_C$ is the fluorescence interior to the oocyte far from the cortex (Fig. S1d). From these measurements, the relative cortical density (RCD) was extracted, using the relation,

$$RCD = \frac{I_0}{I_C - I_{BG}} \tag{8}$$

From each of the two linescans (Fig. S1 b), an RCD value is calculated and the average of these two values is reported as the oocyte's RCD.

#### 2 Active Surface Deformation Model

The first step in interpreting our experimental results is to link the deformation of the oocyte shape to an effective active stress provided by the travelling Rho-band. In various studies this has been achieved within a Hamiltonian model for elastic surfaces [3, 5]. Assuming a travelling Gaussian form for the active stress, the minimization of the Hamiltonian at each time step leads to the expected deformed shape. Meanwhile, several recent studies have explored the deformation of active fluid surfaces under the approximation of thin viscous shells [6–9]. Within this framework the active stress induces velocity flows, which finally cause the cell to deform. Since our ultimate goal is to link the cell deformation to microscopic properties of the actomyosin cortex within an active gel theory [10], we choose for the description of the cortex deformation the latter framework of thin viscous active shells. In order to obtain a more tractable model we make the simplifying assumptions of a fixed cortical thickness and stretching-dominated deformations (membrane limit) [8]. We also examine the deformation in two dimensions (2D), compatible with the 2D experimental data for the outer contour of the oocyte and allow for a friction in the tangential flow, similarly to [11]. In the following, we derive the equations for the active stress induced cell deformation within a purely viscous theory and use them to compare with our experimental data.

##### 2.1 Deformation rate in a thin viscous shell

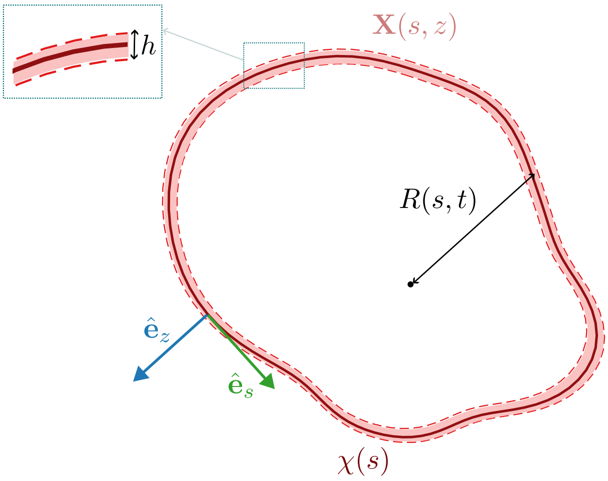

Figure S2: Schematic cartoon depicting coordinates in a cell with a thin actomyosin shell, denoted by a thin pink line.

In this section we derive equations for the rate of deformation of an arbitrarily shaped cell, driven by a thin cortical layer (Fig. S2). For this we solve the force balance $\nabla \cdot \Sigma = 0$ in a thin shell. We start by assuming the following form for the stress tensor

$$\mathbf{\Sigma} = \frac{\eta_s}{2}\left[\nabla \mathbf{v} + (\nabla \mathbf{v})^T\right] + (\eta_b - \eta_s)\mathrm{Tr}\,(\nabla \mathbf{v})\,\mathcal{I} + \Sigma^A \mathcal{I} + \mathcal{T}. \tag{9}$$

This implies that the cortex is a viscous material driven by an active pressure that is generated by its internal mechanics. Here $\eta_b$ and $\eta_s$ are the bulk and shear viscosity, respectively, and $\Sigma^A$ is the magnitude of the active pressure. Furthermore $\mathcal{T}$ is a stress arising from surface tension that acts along the tangential directions of the thin shell.

Given a midsurface curve $\chi(s)$ parameterized by its arclength $s$, we can define its tangent vector $\hat{\mathbf{e}}_s$ and its normal vector $\hat{\mathbf{e}}_z$ as shown in Fig. S2. In the curve-related coordinates $s$ and $z$ a position vector $\mathbf{X}$ can be written as

$$\mathbf{X}(s,z) = \chi(s) + z\hat{\mathbf{e}}_z \tag{10}$$

Note that here, for simplicity, we exploit axisymmetry and thus reduce our description to an effectively two dimensional one. Within the thin shell approximation we assume for the cortex that $z \in [-h/2, h/2]$.
We next introduce the covariant vectors

$$\begin{aligned} \mathbf{e}_s &= \partial_s \mathbf{X} = [1+\kappa(s)z]\hat{\mathbf{e}}_s & (11) \\ \mathbf{e}_z &= \partial_z \mathbf{X} = \hat{\mathbf{e}}_z & (12) \end{aligned}$$

where $\kappa(s) = \left|\frac{d^2\chi}{ds^2}\right|$ is the curvature of the curve $\chi(s)$. Given this we can define the metric tensor

$$g^{ij} = \begin{pmatrix} \frac{1}{[1+\kappa(s)z]^2} & 0 \\ 0 & 1 \end{pmatrix}, \tag{13}$$

which can be used to transform between covariant and contravariant quantities $\mathbf{e}^i = g^{ij}\mathbf{e}_j$. Using this, we write the deformation rate tensor as

$$\begin{aligned} \nabla \mathbf{v} &= \mathbf{e}^i \partial_i v^k \hat{\mathbf{e}}_k \\ &= \hat{\mathbf{e}}_z \hat{\mathbf{e}}_z \partial_z v^z + \hat{\mathbf{e}}_z \hat{\mathbf{e}}_s \partial_z v^s \\ &+ \hat{\mathbf{e}}_s \hat{\mathbf{e}}_z \frac{\partial_s v^z - \kappa(s) v^s}{1+\kappa(s)z} + \hat{\mathbf{e}}_s \hat{\mathbf{e}}_s \frac{\partial_s v^s + \kappa(s) v^z}{1+\kappa(s)z}. \end{aligned} \tag{14}$$

Furthermore, its trace is given by

$$\mathrm{Tr}\,(\nabla \mathbf{v}) = \partial_z v^z + \frac{\partial_s v^s + \kappa(s) v^z}{1+\kappa(s)z}. \tag{15}$$

We can then calculate the expressions for the components the stress tensor using Eqn. 9. These read

$$\begin{aligned} \Sigma^{zz} &= (\eta_b - \eta_s)\frac{\partial_s v^s + \kappa(s) v^z}{1+\kappa(s)z} + \eta_b \partial_z v_z + \Sigma^A \\ \Sigma^{zs} &= \Sigma^{sz} = \frac{\eta_s}{2}\frac{\partial_s v^z - \kappa(s) v^s}{1+\kappa(s)z} + \frac{\eta_s}{2}\partial_z v^s \\ \Sigma^{ss} &= \eta_b \frac{\partial_s v^s + \kappa(s) v^z}{1+\kappa(s)z} + (\eta_b - \eta_s)\,\partial_z v_z + \Sigma^A + \frac{T}{h}, \end{aligned} \tag{16}$$

where we have used $\mathcal{T} = (T/h)\,\hat{\mathbf{e}}_s \hat{\mathbf{e}}_s$, since the surface tension strength can be assumed to act along the tangential direction.
We next seek to rewrite the cortical force balance in the curvilinear coordinates $(s, z)$. In these coordinates, the force density $\nabla \cdot \mathbf{\Sigma}$ is given by

$$\begin{aligned} \nabla \cdot \mathbf{\Sigma} &= \mathbf{e}^i \cdot \partial_i \Sigma^{kj} \hat{\mathbf{e}}_k \hat{\mathbf{e}}_j \\ &= \hat{\mathbf{e}}_s \left\{ \frac{1}{1+\kappa(s)z}\partial_s \Sigma^{ss} + \partial_z \Sigma^{zs} + \frac{2\kappa(s)}{1+\kappa(s)z}\Sigma^{zs} \right\} \\ &+ \hat{\mathbf{e}}_z \left\{ \frac{1}{1+\kappa(s)z}\partial_s \Sigma^{sz} + \partial_z \Sigma^{zz} - \frac{\kappa(s)}{1+\kappa(s)z}\left(\Sigma^{ss} - \Sigma^{zz}\right) \right\}. \end{aligned} \tag{17}$$

Thus, putting everything together, in the absence of external forces the force balance, $\boldsymbol{\nabla} \cdot \mathbf{\Sigma} = \mathbf{0}$, in the tangential direction $s$ reads

$$\partial_s \Sigma^{ss} + \partial_z \left\{ [1+\kappa(s)z]\Sigma^{zs} \right\} + \kappa(s)\Sigma^{zs} = 0, \tag{18}$$

and the force balance in the normal direction $z$, reads

$$\partial_s \Sigma^{sz} + \partial_z \left\{ [1+\kappa(s)z]\,\Sigma^{zz} \right\} - \kappa(s)\Sigma^{ss} = 0 \tag{19}$$

with the respective components of the stress provided in Eqs. 16.

We next exploit the fact that the cell cortex is thin, i.e., $h \ll R(s)$, assume that the cortical thickness is conserved at a constant value $h$ and integrate over the normal direction $z$ from $-h/2$ to $h/2$, keeping only the first order in $h/R$. Under this approximation we have for the components $\Sigma^{ij}(s,z)$, provided in Eqs. 16, that

$$\int_{-h/2}^{h/2} \Sigma^{ij}(s,z)dz = h\Sigma^{ij}(s,0) + \mathcal{O}(h^2/R^2). \tag{20}$$

Along these lines we introduce the averaged cortical velocities in the $s$ and $z$ direction as $V^s = v^s(s,0)$ and $V^z = v^z(s,0)$. We also assume incompressibility of the cortex, implying that

$$\partial_z v^z|_{z=0} = -\frac{\partial_s v^s + \kappa(s)v^z}{1+\kappa(s)z}\bigg|_{z=0}. \tag{21}$$

For the boundary conditions in the tangential direction we assume

$$\Sigma^{zs}(s,h/2) = -\gamma h V^s \text{ and } \Sigma^{zs}(s,-h/2) = 0, \tag{22}$$

where $\gamma$ denotes the friction coefficient between the cortex and the surrounding membrane. These conditions imply that at first order in the thickness $h$ we have $\Sigma^{zs}(s,0) = 0$ and

$$\int_{-h/2}^{h/2} \partial_z \left\{[1+\kappa(s)z]\Sigma^{zs}\right\} dz = -\gamma h V^s. \tag{23}$$

In the normal direction we assume that

$$\Sigma^{zz}(s,h/2) = \Delta P \text{ and } \Sigma^{zz}(s,-h/2) = 0, \tag{24}$$

with $\Delta P$ denoting the pressure difference between the inside and the outside of the cell leading to

$$\int_{-h/2}^{h/2} \partial_z \left\{[1+\kappa(s)z]\Sigma^{zz}\right\} dz = \Delta P\left(1+\frac{\kappa h}{2}\right). \tag{25}$$

For simplicity we also assume here that the cortical viscosities $\eta_s$ and $\eta_b$ are height independent. Under the above assumptions (Eqs. (20), (21), (22), (23), (24), (25)) the thin-shell approximation leads to the following force balance equations in the $s$ direction

$$\eta_s h\partial_s\left(\partial_s V^s + \kappa V^z\right) + h\partial_s\Sigma^A - \gamma h V_s = 0 \tag{26}$$

and the $z$ direction respectively

$$\kappa\eta_s h\left(\partial_s V^s + \kappa V^z\right) + \kappa h\Sigma^A + \kappa T - \Delta P\left(1+\frac{\kappa h}{2}\right) = 0. \tag{27}$$

In steady state $V^s, V^z$ are zero and the Laplace equation $\kappa T - \Delta P\left(1+\frac{\kappa h}{2}\right) = 0$ holds, leading to a circular shape with a constant radius $R$ and curvature $\kappa = \frac{1}{R}$. The $z$-direction in this case coincides with the radial direction $r$ and the arc length coordinate is proportional to the angle $\theta$, namely $s = R\theta$. The linearized equations of motion around this state thus become

$$\frac{\eta_s}{R^2}\left(\partial_\theta^2 V^s + \partial_\theta V^z\right) + \frac{1}{R}\partial_\theta\Sigma^A - \gamma V^s = 0 \tag{28}$$

and

$$\frac{\eta_s}{R^2}\left(\partial_\theta V^s + V^z\right) + \frac{1}{R}\Sigma^A = 0. \tag{29}$$

We next Fourier transform in the $\theta$ direction and solve for $\tilde{V}^z = \int_{-\infty}^{+\infty} dq\ V^z \exp(iq\theta)$ to obtain the following expression for radial velocity

$$\frac{\tilde{V}^z}{R} = -\frac{\tilde{\Sigma}^A}{\eta_s}, \tag{30}$$

implying the rate of deformation of the oocyte is expected to be proportional to $\Sigma^A/\eta$, which is what we use in the main text.

#### 2.2 Predictions of the thin viscous shell theory for the cell dynamics

We have seen how the force balance equations (26) and (27) determine the midsurface tangential velocity $V^s$ and normal velocity $V^z$ for a given oocyte shape, active stress and pressure. In turn, the normal velocity $V^z$ induces a change of shape in time, while $V^s$ gives only a change in frame for the parametrization of the shell [12]. Since we are interested only in the characterization of the oocyte dynamics we focus here on the effect of $V^z$.
Given the angular parametrization of the midsurface curve (Fig. S2)

$$\chi(\theta, t) = R(\theta, t)\cos\theta\ \hat{\mathbf{e}}_x + R(\theta, t)\sin\theta\ \hat{\mathbf{e}}_y \tag{31}$$

we obtain

$$V^z = \frac{R}{S}\frac{dR}{dt} \tag{32}$$

and

$$V^s = \frac{\partial_\theta R}{S}\frac{dR}{dt} \tag{33}$$

where

$$S(\theta, t) = \frac{ds}{d\theta} = |\partial_\theta \chi(\theta, t)| = \sqrt{R^2 + (\partial_\theta R)^2}. \tag{34}$$

Within this angular parametrization the force balance equations read

$$\frac{1}{S}\partial_\theta\left(\frac{1}{S}\partial_\theta V^s + \kappa V^z\right) + \frac{1}{S}\partial_\theta\frac{\Sigma^A}{\eta_s} - \frac{\gamma}{\eta_s}V_s \quad = \quad 0 \tag{35}$$

and

$$\frac{\kappa}{S}\partial_\theta V^s + \kappa^2 V^z + \kappa\frac{\Sigma^A}{\eta_s} + \frac{1}{h\eta_s}\left[\kappa T - \Delta P\left(1 + \frac{\kappa h}{2}\right)\right] = 0. \tag{36}$$

Together Eqs. 35, 36 and 32 can determine the dynamics of the cell deformation during the SCW within the viscous shell theory. We can use these in two ways. First, given the experimental data for the cell shape deformation in terms of the local radius $R(\theta, t)$ we can estimate the active stress $\Sigma^A(\theta, t)/\eta_s$ of the SCW. Second assuming a particular $\Sigma^A(\theta, t)/\eta_s$ profile we can numerically solve the set of equations to extract the predicted by the viscous theory shape deformation $R(\theta, t)$. A comparison of the latter with the experimental one can be used to judge the validity of the theory, as discussed in Figure 6 of the main text. We remark that in all cases we assume that the viscosity is constant within the cortex and define all the quantities in terms of it.

##### 2.2.1 Active stress estimation from experimental data

Starting by the local radial profile $R(\theta, t)$ we can calculate the normal velocity $V^z(\theta, t)$ from Eq. 32. Then we can estimate the tangential velocity $V^s(\theta, t)$ from Eq. 33. Assuming a small value for the friction parameter $\gamma/\eta_s = 0.0001\mu\text{m}^{-2}$, consistent with the findings of [11], we can estimate the active stress $\Sigma^A(\theta, t)/\eta_s$ from Eq. 35. Note that the latter equation provides us with $\Sigma^A(\theta, t)/\eta_s$ up to a constant $C(t)$, which we choose it so that it is positive for all $\theta, t$. This makes up for the balance in pressure as dictated by Eq. 36. Our results are shown for the control case in Fig. S3.

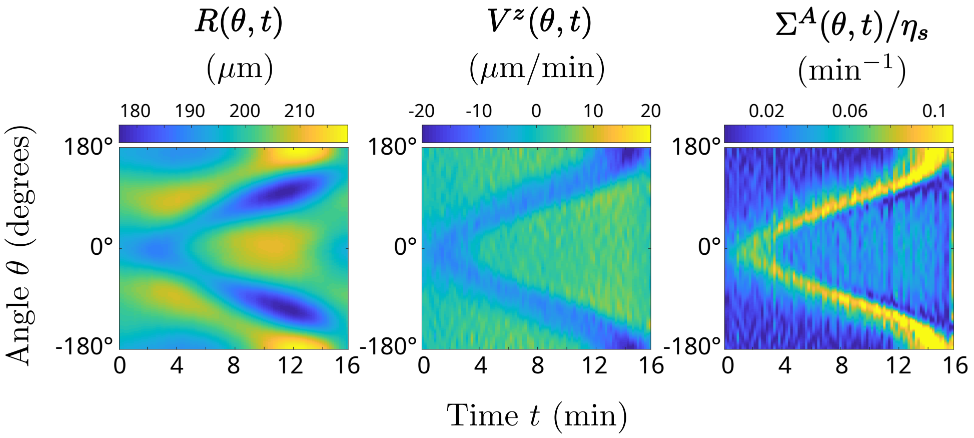

Figure S3: Calculation of resulting radial profile $R(\theta, t)$ for a given active stress $\Sigma^A(\theta, t)/\eta_s$ of the form of Eq. 37 with a value $B = 0.114$. As an intermediate step the normal velocity $V^z(\theta, t)$ should be estimated. The radial profile $R(\theta, t)$ is obtained after shifting the center of mass of the calculated radius at each time step, so that it is always centered around $(0, 0)$(see Sec. 2.2.3). All the parameter values used are shown in table 1.

##### 2.2.2 Numerical prediction of cell deformation for assumed active stress

In order to verify our viscous theory we follow here the opposite procedure. We start with assuming an active stress profile

$$\Sigma^A(\theta,t)/\eta_s = B\frac{\exp\left[w\cos\left(\theta-\omega t\right)\right]+\exp\left[w\cos\left(\theta+\omega t\right)\right]}{\max_\theta\left\{\exp\left[w\cos\left(\theta-\omega t\right)\right]+\exp\left[w\cos\left(\theta+\omega t\right)\right]\right\}} \tag{37}$$

in the form of a propagating circular Gaussian (von-Mises) distribution with a chosen maximum strength $B$, a width regulating parameter $w = 8$ and an angular velocity $\omega = 0.196$rad/min and use it as an input for predicting the cell deformation through equations (35), (36) and (32). Both the chosen profile and the selected parameters were found to match reasonably well the active stress profiles extracted in experiments for suitable values of $B$ (compare Figs. S3 and S4, as well as Fig. 3, main text) . As a way to account for the volume of the cell being approximately constant and our lack of knowledge of the experimentally applied pressure, we replace the Laplace law in Eq. (36) with a Lagrange multiplier $p$ to be determined, similarly to [6]. This leads to the following expression for the force balance in the $z$ direction

$$\frac{\kappa}{S}\partial_\theta V^s + \kappa^2 V^z + \kappa\frac{\Sigma^A}{\eta_s} + \frac{1}{h\eta_s}p + \frac{\gamma_z}{\eta_s}V^z = 0. \tag{38}$$

with the constant volume constraint, corresponding here to a constant area $A$ inside the curve, i.e. $dA/dt = 0$, provided by

$$\int_0^{2\pi} d\theta\ V^z S = 0. \tag{39}$$

Note that here we have also introduced a very small friction $\gamma_z/\eta_s = 10^{-9}\mu\text{m}^{-2}$ in the $z$ direction to enhance numerical stability.

Solving numerically Eqs. (35),(38), (39) and (32) we are able to calculate the cell deformation. More specifically, we introduce an $N$-point grid $\theta_i = \frac{2\pi i}{N}$. From the cell shape at time $t_j$, $R(\theta_i, t_j)$ we can calculate the corresponding curvature $\kappa$ and arc-length $S$. Given $\Sigma^A(\theta_i, t_j)/\eta_s$,

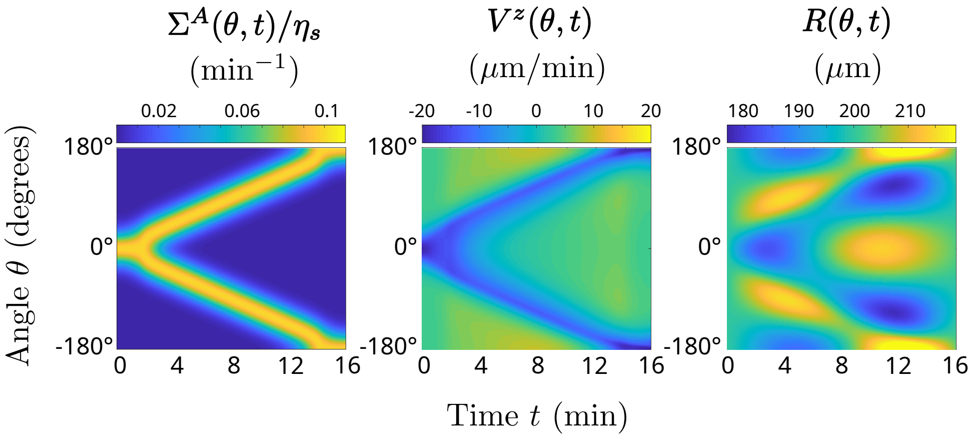

Figure S4: Computation of active stress profile $\Sigma^A(\theta,t)/\eta_s$ from the experimental radial profile $R(\theta,t)$ for the control case. As an intermediate step the normal velocity $V^z(\theta,t)$ should be estimated from Eq.32, after recovering the center-of-mass motion (see Sec. 2.2.3)

we calculate the velocities $V^z(\theta_i,t_j), V^s(\theta_i,t_j)$ and pressure $p(t_j)$, solving numerically the non-linear system of equations 35, 38 and 39 on the grid. We can then obtain the cell shape at time $t_{j+1}=t_j+dt$, $R(\theta_i,t_{j+1})$, by numerically integrating Eq. 32, i.e

$$\frac{dR}{dt}=\frac{S(R)}{R}V_z \tag{40}$$

with an implicit-explicit Euler method for enhanced stability. In particular, we subtract in both sides a diffusion term $\alpha\partial_\theta^2 R$ from Eq. 32 and solve one part implicitly $(im)$ and the other part explicitly $(ex)$

$$\left[\frac{dR}{dt}-\alpha\partial_\theta^2 R\right]^{(im)}=\left[\frac{S(R)}{R}V_z-\alpha\partial_\theta^2 R\right]^{(ex)} \tag{41}$$

using the Euler method. An iteration over the whole duration of the SCW provides us with the corresponding cell deformation $R(\theta,t)$. For our numerics we start from a circular cell shape $R(\theta,0)=R_0(\cos\theta,\sin\theta)$, with $R_0=200\mu$m. We use a grid size $N=256$, a step size $dt=0.02$min, a friction coefficient $\gamma/\eta_s=0.0001\mu\text{m}^{-2}$, $\alpha=1.25\text{min}^{-1}$ and integrate for $N_t=800$ steps. All our spatial derivatives are calculated with periodic two-point central difference. All the simulation parameters are summarized in Table 1 and a typical simulation result corresponding to the wild type case of Fig. S3 is shown in Fig. S4. We have repeated our numerics for smaller step-sizes and values $\alpha$ and found that it leads to similar results. We nevertheless note that for choosing our parameters we prioritized stability over numerical accuracy. A higher level of accuracy an be possibly reached by employing more sophisticated methods as in [6, 12].

##### 2.2.3 A note on the center of mass motion

As mentioned in Sec 1.4.1 in order to estimate the deformation rate from experimental oocyte images we fit the $x$,$y$ coordinates of the oocyte shapes to circles, extract their center and choose the polar coordinates of our description so that $r=0$ always correspond to the center of the fitted circle. This roughly amounts to shifting the center of mass of the oocyte shapes so that it always lies at $(0,0)$. Although this is advantageous when comparing different shapes it hides the fact that the center of mass inevitably moves in response to a shape change (Fig. S5, 3rd panel). Since within our theoretical framework we assume a fixed coordinate space the radial deformation rate $\frac{dR}{dt}$ includes the center of mass motion. Thus, when we try to estimate the

**Simulation Parameters**

| | | | |
|---|---|---|---|
| Angular velocity of active stress $(\Sigma_A/\eta_s)$ strength | $\omega$ | $[\Theta/T]$ | 0.196 rad/min |
| Active stress $(\Sigma_A/\eta_s)$ width regulating parameter | $\alpha$ | [1] | 8 |
| Active stress $(\Sigma_A/\eta_s)$ strength | $B$ | $[1/T]$ | $0.007 - 0.158\text{min}^{-1}$ |
| Initial oocyte radius | $R_0$ | $[L]$ | $200\mu$m |
| Cortex width | $h$ | $[L]$ | $10\mu$m |
| Friction coefficient (tangential) | $\gamma/\eta_s$ | $[1/L^2]$ | $0.0001\mu\text{m}^{-2}$ |
| Friction coefficient (normal) | $\gamma_z/\eta_s$ | $[1/L^2]$ | $10^{-9}\mu\text{m}^{-2}$ |
| Number of spatial grid points | $N$ | [1] | 256 |
| Number of time steps | $N_t$ | [1] | 800 |
| Time step | $dt$ | $[T]$ | 0.02min |
| Diffusion parameter | $\alpha$ | $[1/T]$ | $1.25\text{min}^{-1}$ |

Table 1: Summary of the simulation parameters for the numerical prediction of cell deformation for assumed active stress. The third column shows the units of the parameters in terms of arbitrary angle $\Theta$,length $L$ and time $T$ units.

active stress from experimental data we need first to recover the radial profiles with the center of mass motion (Fig. S5, 2nd panel), calculate then $\frac{dR}{dt}$ and from this derive the active stress $\Sigma^A$. Similarly, when we estimate numerically the characteristic deformation rate, we need to follow the experimental procedure and shift all radial profiles so that they are centered around $(0,0)$, so that we can calculate the local deformation rate $d(\theta,t)$ (Eq. (2)).

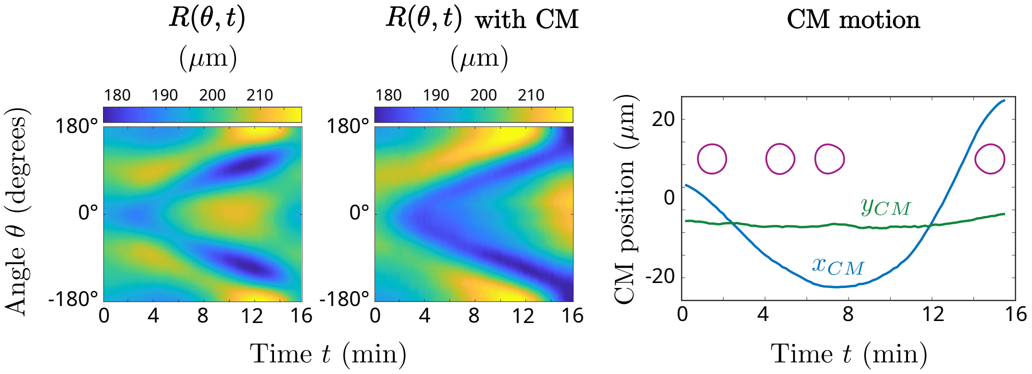

Figure S5: Effect of the center of mass (CM) motion on the radial kymograph profile $R(\theta,t)$ from experimental data for the control case. The actual center-of-mass motion is shown in the third panel, which depicts also the corresponding oocyte shapes for selected times.

### 3 Active Fluid Model

#### 3.1 Continuum description of an isotropic actomyosin gel

In this section we derive the stress exerted by passive and motor crosslinkers in an isotropic actomyosin gel. Our derivation closely follows the logic outlined in [10].

##### 3.1.1 Continuous Field variable

We model the cell cortex as consisting of actin polymer filaments, passive crosslinkers, and molecular-scale motors. In our theory, the $i$-th filament is characterized by its center of mass

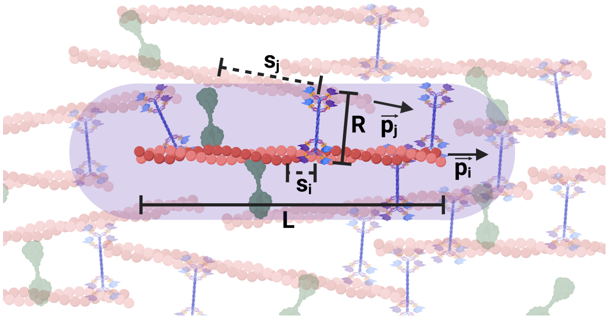

Figure S6: Parameter definitions for interactions between actin filaments (red) mediated by molecular motors (blue) and passive crosslinkers (green).

position $\mathbf{x}_i$ and the direction in which it points $\mathbf{p}_i$, where $\mathbf{p}_i$ is a unit vector (Fig. S6). For simplicity we take all filaments to have the same length $L$.

Since we are interested in phenomena that occur on length scales that are large compared to individual filaments ($R_{cell} \sim \mathcal{O}(100\ \mu m)$, $L_{filament} \sim \mathcal{O}(1\ \mu m)$ ), we seek to describe the large scale physics of the cell cortex, in terms of continuous fields. We thus define the filament mass density $\rho$ as

$$\rho(\mathbf{x}) = L \sum_i \delta(\mathbf{x}_i - \mathbf{x}). \tag{42}$$

and the filament center of mass velocity $\mathbf{v}$ as,

$$\rho(\mathbf{x})\mathbf{v}(\mathbf{x}) = L \sum_i \mathbf{v}_i \delta(\mathbf{x}_i - \mathbf{x}). \tag{43}$$

In principle, actin filaments in the cortex could also be directionally ordered. Here we take the cortex to be isotropic, and thus the polar order $\mathbf{P}$, which is defined by

$$\rho(\mathbf{x})\mathbf{P}(\mathbf{x}) = L \sum_i \mathbf{p}_i \delta(\mathbf{x}_i - \mathbf{x}), \tag{44}$$

vanishes and $\mathbf{P} = 0$. Additionally, the nematic order tensor $\mathcal{Q}$, defined by

$$\rho(\mathbf{x})\mathcal{Q}(\mathbf{x}) = L \sum_i \mathbf{p}_i \mathbf{p}_i \delta(\mathbf{x}_i - \mathbf{x}), \tag{45}$$

obeys $\mathcal{Q} = \mathcal{I}/3$, where $\mathcal{I}$ is the identity tensor.

For notational convenience, we also introduce the tensors $\mathcal{H}$ and $\mathcal{J}$, defined as,

$$\rho(\mathbf{x})\mathcal{H}(\mathbf{x}) = L \sum_i \mathbf{p}_i \dot{\mathbf{p}}_i \delta(\mathbf{x}_i - \mathbf{x}) \tag{46}$$

and

$$\rho(\mathbf{x})\mathcal{J}(\mathbf{x}) = L \sum_i \mathbf{p}_i \left(\mathbf{v}_i - \mathbf{v}(\mathbf{x})\right) \delta(\mathbf{x}_i - \mathbf{x}). \tag{47}$$

##### 3.1.2 Modelling motorized and passive crosslinks

We start by considering the system on long timescales and postulate the force density $\mathbf{f}_{ij}$ exerted by motors connecting two filaments labelled $i$ and $j$ as,

$$\mathbf{f}_{ij}^M = -\gamma^M c^M(s_i, s_j) \left(\mathbf{v}_i + s_i \dot{\mathbf{p}}_i - \mathbf{v}_j - s_j \dot{\mathbf{p}}_j + V_{||}(\mathbf{p}_i - \mathbf{p}_j)\right), \tag{48}$$

where $c^M(s_i, s_j)$ is the density of motors bound between point $s_i$ on filament $i$ and point $s_j$ on filament $j$. Furthermore, $V_{||}$ is the unloaded speed of the motor, and $-\gamma_M$ is the slope of the motor's force-velocity curve. This means that if $V_{||} > 0$, the motor walks in a direction parallel to $\mathbf{p}_i$. We also define the arc-length coordinate $s_i$, such that $s_i = L/2$ denotes the plus end of the filament.

Similarly, we take the passive crosslinker density to obey

$$\mathbf{f}_{ij}^X = -\gamma^X c^X(s_i, s_j)\left(\mathbf{v}_i + s_i\dot{\mathbf{p}}_i - \mathbf{v}_j - s_j\dot{\mathbf{p}}_j\right) \tag{49}$$

In order to respect the invariance of the system to Galilean tranformations and global rotations we have to project the above expressions for the force densities along the orientation of the $ij$ (motorized or passive) crosslinker, provided by

$$\hat{\mathbf{w}}_{ij} = \frac{\mathbf{x}_i + s_i\mathbf{p}_i - \mathbf{x}_j - s_j\mathbf{p}_j}{|\mathbf{x}_i + s_i\mathbf{p}_i - \mathbf{x}_j - s_j\mathbf{p}_j|}. \tag{50}$$

Thus, the expressions for the motor and passive crosslinker force densities are provided by

$$\mathbf{f}_{ij}^{M(X)} = \left(\mathbf{f}_{ij}^{M(X),0} \cdot \hat{\mathbf{w}}_{ij}\right)\hat{\mathbf{w}}_{ij}. \tag{51}$$

We next postulate the functional forms for the motor and crosslinker densities, given by,

$$c^M(s_i, s_j) = C_0^M + (s_i + s_j)C_1^M \tag{52}$$

and

$$c^X(s_i, s_j) = C_0^X + (s_i + s_j)C_1^X \tag{53}$$

for the densities of doubly-bound motors and passive crosslinkers, respectively. Note that in general terms which are higher order in $s_i, s_j$ are allowed. However, they do not not contribute to the stress tensor [10], and thus we ignore them here for simplicity.

Finally crosslinks have a characteristic size $R$, which is the maximum range over which they can connect two filaments. Here, we take $R$ to be the same for motors and passive crosslinkers, for simplicity.

##### 3.1.3 Force Balance on single filaments

In overdamped systems, such as the cell cortex, the total force on each filament vanishes, $\mathbf{F}_i = 0$. Following reference [10], we write

$$\mathbf{F}_i = \sum_j \int_{-L/2}^{L/2} ds_i \int_{-L/2}^{L/2} ds_j \int_{\Omega(\mathbf{x}_i)} d^3\mathbf{x}'\delta\left(\mathbf{x}' - \mathbf{x}_j - s_j\mathbf{p}_j + s_i\mathbf{p}_i\right)\left[\left(\mathbf{f}_{ij}^{M,0} + \mathbf{f}_{ij}^{X,0}\right)\cdot\hat{\mathbf{w}}_{ij}\right]\hat{\mathbf{w}}_{ij} \tag{54}$$

where $\Omega(\mathbf{x}_i)$ denotes a spherical domain of radius $R$ centered at $\mathbf{x}_i$.
The lowest order terms read (see also [5])

$$\mathbf{F}_i = \sum_j \int_{-L/2}^{L/2} ds_i \int_{-L/2}^{L/2} ds_j \int_{\Omega(\mathbf{x}_i)} d^3\mathbf{x}'\delta\left(\mathbf{x}' - \mathbf{x}_j\right)\left[\left(\mathbf{f}_{ij}^{M,0} + \mathbf{f}_{ij}^{X,0}\right)\cdot\left(\mathbf{x}_i - \mathbf{x}'\right)\right]\frac{\mathbf{x}_i - \mathbf{x}'}{|\mathbf{x}_i - \mathbf{x}'|^2} + \mathcal{O}\left(R^3L^3\right) \tag{55}$$

After evaluating the integrals over $s_i, s_j$ and introducing

$$\mathcal{A}\left(\mathbf{x}_i - \mathbf{x}'\right) = \frac{\left(\mathbf{x}_i - \mathbf{x}'\right)\left(\mathbf{x}_i - \mathbf{x}'\right)}{|\mathbf{x}_i - \mathbf{x}'|^2} \tag{56}$$

we obtain in lowest order,

$$\begin{aligned}\mathbf{F}_i = &-L^2\left(C_0^M\gamma^M + C_0^X\gamma^X\right)\sum_j\int_{\Omega(\mathbf{x}_i)} d^3\mathbf{x}'\delta\left(\mathbf{x}'-\mathbf{x}_j\right)\left[\left(\mathbf{v}_i-\mathbf{v}_j\right)\cdot\mathcal{A}\left(\mathbf{x}_i-\mathbf{x}'\right)\right]\\ &- L^2C_0^M\gamma^M V_{||}\sum_j\int_{\Omega(\mathbf{x}_i)} d^3\mathbf{x}'\delta\left(\mathbf{x}'-\mathbf{x}_j\right)\left[\left(\mathbf{p}_i-\mathbf{p}_j\right)\cdot\mathcal{A}\left(\mathbf{x}_i-\mathbf{x}'\right)\right]+\mathcal{O}\left(R^3L^3\right)\end{aligned} \tag{57}$$

Next, executing the $j$ sum, using the definitions Eqns. 28, 29, 30 leads to

$$\begin{aligned}\mathbf{F}_i = &-L\left(C_0^M\gamma^M + C_0^X\gamma^X\right)\int_{\Omega(\mathbf{x}_i)} d^3\mathbf{x}'\left[\left(\mathbf{v}_i-\mathbf{v}\left(\mathbf{x}'\right)\right)\cdot\mathcal{A}\left(\mathbf{x}_i-\mathbf{x}'\right)\right]\\ &- L^2C_0^M\gamma^M V_{||}\int_{\Omega(\mathbf{x}_i)} d^3\mathbf{x}'\left[\mathbf{p}_i\cdot\mathcal{A}\left(\mathbf{x}_i-\mathbf{x}'\right)\right]+\mathcal{O}\left(R^3L^3\right)\end{aligned} \tag{58}$$

Finally, we Taylor expand $\rho\left(\mathbf{x}'\right)=\rho\left(\mathbf{x}_i\right)+\mathcal{O}(R)$ and $\mathbf{v}\left(\mathbf{x}'\right)=\mathbf{v}\left(\mathbf{x}_i\right)+\mathcal{O}(R)$ and do the volume integral

$$\int_{\Omega(\mathbf{x}_i)} d^3\mathbf{x}'\mathcal{A}\left(\mathbf{x}_i-\mathbf{x}'\right)=\frac{4\pi R^3}{9}\mathcal{I} \tag{59}$$

to arrive at

$$\mathbf{F}_i=-\frac{4\pi R^3L}{9}\rho(\mathbf{x}_i)\left\{\left(C_0^M\gamma^M+C_0^X\gamma^X\right)\left(\mathbf{v}_i-\mathbf{v}\left(\mathbf{x}_i\right)\right)+C_0^M\gamma^M V_{||}\mathbf{p}_i\right\}+\mathcal{O}\left(R^3L^3\right) \tag{60}$$

and thus, since $\mathbf{F}_i=\mathbf{0}$,

$$\mathbf{v}_i-\mathbf{v}=-V_{||}\frac{C_0^M\gamma^M}{C_0^M\gamma^M+C_0^X\gamma^X}\mathbf{p}_i+\mathcal{O}(L^2). \tag{61}$$

It directly follows that

$$\mathcal{J}=-\frac{V_{||}}{3}\frac{C_0^M\gamma^M}{C_0^M\gamma^M+C_0^X\gamma^X}\mathcal{I}+\mathcal{O}(L^2). \tag{62}$$

##### 3.1.4 Torque Balance on single filaments

In the overdamped system that we consider here, the total torque on each filament vanishes, $\mathbf{T}_i=0$. The torque balance on filament $i$ implies that

$$\begin{aligned}\mathbf{T}_i = &\sum_j\int_{-L/2}^{L/2}ds_i\int_{-L/2}^{L/2}ds_j\int_{\Omega(\mathbf{x}_i)}d^3\mathbf{x}'\delta\left(\mathbf{x}'-\mathbf{x}_j\right)\left\{s_i\mathbf{p}_i\times\left[\left(\mathbf{f}_{ij}^{M,0}+\mathbf{f}_{ij}^{X,0}\right)\cdot\mathcal{A}\left(\mathbf{x}_i-\mathbf{x}'\right)+\boldsymbol{\tau}_{ij}\right]\right\}\\ &+\sum_j\int_{-L/2}^{L/2}ds_i\int_{-L/2}^{L/2}ds_j\int_{\Omega(\mathbf{x}_i)}d^3\mathbf{x}'\left\{\left(s_i\mathbf{p}_i-s_j\mathbf{p}_j\right)\cdot\nabla'\delta\left(\mathbf{x}'-\mathbf{x}_j\right)\right\}\left\{s_i\mathbf{p}_i\times\left[\left(\mathbf{f}_{ij}^{M,0}+\mathbf{f}_{ij}^{X,0}\right)\cdot\mathcal{A}\left(\mathbf{x}_i-\mathbf{x}'\right)\right]\right\}\\ &+\sum_j\int_{-L/2}^{L/2}ds_i\int_{-L/2}^{L/2}ds_j\int_{\Omega(\mathbf{x}_i)}d^3\mathbf{x}'\left\{\left(s_i\mathbf{p}_i-s_j\mathbf{p}_j\right)\cdot\nabla'\delta\left(\mathbf{x}'-\mathbf{x}_j\right)\right\}\left\{s_i\mathbf{p}_i\times\boldsymbol{\tau}_{ij}\right\}\end{aligned} \tag{63}$$

where $\tau_{ij}$ results is the explicit torque that crosslinks exert between filaments $i$ and $j$. Such a torque could for instance result from crosslinks acting as torsional springs. Following reference [10], here we set $\tau_{ij}=0$ for simplicity.

We now write

$$\mathbf{T}_i = \sum_j \int_{-L/2}^{L/2} ds_i \int_{-L/2}^{L/2} ds_j \int_{\Omega(\mathbf{x}_i)} d^3\mathbf{x}'\ s_i\mathbf{p}_i \times \left[\left(\mathbf{f}_{ij}^{M,0} + \mathbf{f}_{ij}^{X,0}\right) \cdot \left(\mathcal{A}\left(\mathbf{x}_i - \mathbf{x}'\right)\delta\left(\mathbf{x}' - \mathbf{x}_j\right)\right)\right]$$
$$+ \sum_j \int_{-L/2}^{L/2} ds_i \int_{-L/2}^{L/2} ds_j \int_{\Omega(\mathbf{x}_i)} d^3\mathbf{x}'\ s_i^2\mathbf{p}_i \times \left(\mathbf{p}_i\cdot\nabla'\right)\left[\left(\mathbf{f}_{ij}^{M,0} + \mathbf{f}_{ij}^{X,0}\right) \cdot \left(\mathcal{A}\left(\mathbf{x}_i - \mathbf{x}'\right)\delta\left(\mathbf{x}' - \mathbf{x}_j\right)\right)\right] \tag{64}$$

Following the same sequence of steps as above, and using Eqn. 61 we arrive at,

$$\dot{\mathbf{p}}_i = \left(\mathcal{I} - \mathbf{p}_i\mathbf{p}_i\right)\cdot\left(\mathbf{p}_i\cdot\nabla\mathbf{v}\right). \tag{65}$$

It follows that $\mathcal{H}$, in the isotropic limit, is given by

$$\mathcal{H} = \frac{1}{3}\nabla\mathbf{v} - \frac{1}{15}\left[\mathrm{Tr}\left(\nabla\mathbf{v}\right)\mathcal{I} + \nabla\mathbf{v} + \left(\nabla\mathbf{v}\right)^T\right]. \tag{66}$$

#### 3.2 Material Stresses

We now calculate the stress exerted by motors using the prescription

$$\mathbf{\Sigma}^M(\mathbf{x}) = -\frac{1}{2}\sum_{ij}\int_{-L/2}^{L/2} ds_i \int_{-L/2}^{L/2} ds_j \int_{\Omega(\mathbf{x}_i)} d^3\mathbf{x}'\delta\left(\mathbf{x}-\mathbf{x}_i\right)\delta\left(\mathbf{x}'-\mathbf{x}_j - s_j\mathbf{p}_j + s_i\mathbf{p}_i\right)\left[\mathbf{x}_i - \mathbf{x}_j\right]\mathbf{f}_{ij}^M \tag{67}$$

Upon Taylor expanding the second $\delta$ function around $\mathbf{x}' - \mathbf{x}_j$ we obtain

$$\mathbf{\Sigma}^M(\mathbf{x}) = \mathbf{\Sigma}^{M,0}(\mathbf{x}) + \mathbf{\Sigma}^{M,1}(\mathbf{x}) + \mathbf{\Sigma}^{M,2}(\mathbf{x}) + \mathcal{O}\left(L^5\right), \tag{68}$$

with

$$\mathbf{\Sigma}^{M,0}(\mathbf{x}) = -\frac{1}{2}\sum_{ij}\int_{-L/2}^{L/2} ds_i \int_{-L/2}^{L/2} ds_j \int_{\Omega(\mathbf{x}_i)} d^3\mathbf{x}'\delta\left(\mathbf{x}-\mathbf{x}_i\right)\delta\left(\mathbf{x}'-\mathbf{x}_j\right)\left[\mathbf{x}_i - \mathbf{x}_j\right]\left[\mathbf{f}_{ij}^{M,0}\cdot\mathcal{A}\left(\mathbf{x}_i - \mathbf{x}'\right)\right], \tag{69}$$

$$\mathbf{\Sigma}^{M,1}(\mathbf{x}) =$$
$$-\frac{1}{2}\sum_{ij}\int_{-L/2}^{L/2} ds_i \int_{-L/2}^{L/2} ds_j \int_{\Omega(\mathbf{x}_i)} d^3\mathbf{x}'\delta\left(\mathbf{x}-\mathbf{x}_i\right)\left[\left(s_i\mathbf{p}_i - s_j\mathbf{p}_j\right)\cdot\nabla'\delta\left(\mathbf{x}'-\mathbf{x}_j\right)\right]\left[\mathbf{x}_i - \mathbf{x}_j\right]\left[\mathbf{f}_{ij}^{M,0}\cdot\mathcal{A}\left(\mathbf{x}_i - \mathbf{x}'\right)\right], \tag{70}$$

and

$$\mathbf{\Sigma}^{M,2}(\mathbf{x}) = -\frac{1}{4}\sum_{ij}\int_{-L/2}^{L/2} ds_i \int_{-L/2}^{L/2} ds_j \int_{\Omega(\mathbf{x}_i)} d^3\mathbf{x}'\delta\left(\mathbf{x}-\mathbf{x}_i\right)\left[\left(s_i^2\mathbf{p}_i\mathbf{p}_i + s_j^2\mathbf{p}_j\mathbf{p}_j\right):\nabla'^2\delta\left(\mathbf{x}'-\mathbf{x}_j\right)\right]\left[\mathbf{x}_i - \mathbf{x}_j\right]$$
$$\left[\mathbf{f}_{ij}^{M,0}\cdot\mathcal{A}\left(\mathbf{x}_i - \mathbf{x}'\right)\right], \tag{71}$$

with the shorthand : denoting the contraction in two indices.
For clarity, we make use of the index notation in the following, employing small greek letters to denote the different coordinates and implying summation over repeated indices. Employing this notation and performing the integrals over $ds_i$ and $ds_j$ gives

$$\mathbf{\Sigma}^{M,0}_{\beta\alpha}(\mathbf{x}) = \frac{\gamma^M}{2} C_0^M L^2 \sum_{ij} \int_{\Omega(\mathbf{x})} d^3\mathbf{x}' \delta\left(\mathbf{x}-\mathbf{x}_i\right) \delta\left(\mathbf{x}'-\mathbf{x}_j\right) \left[\mathbf{x}-\mathbf{x}'\right]_\beta \mathcal{A}_{\alpha\zeta}\left(\mathbf{x}-\mathbf{x}'\right) \left[\mathbf{v}_i - \mathbf{v}_j + V_{||}\left(\mathbf{p}_i - \mathbf{p}_j\right)\right]_\zeta + \mathcal{O}(L^4R^4) \tag{72}$$

$$\begin{aligned}\mathbf{\Sigma}^{M,1}_{\beta\alpha}(\mathbf{x}) = {} & \frac{\gamma^M}{24} C_0^M L^4 \sum_{ij} \int_{\Omega(\mathbf{x})} d^3\mathbf{x}' \mathcal{A}_{\alpha\zeta}\left(\mathbf{x}-\mathbf{x}'\right) \nabla'_\kappa \left\{ \left[\mathbf{x}_i - \mathbf{x}_j\right]_\beta \delta\left(\mathbf{x}-\mathbf{x}_i\right) \delta\left(\mathbf{x}'-\mathbf{x}_j\right) \left[\mathbf{p}_i\dot{\mathbf{p}}_i + \mathbf{p}_j\dot{\mathbf{p}}_j\right]_{\kappa\zeta} \right\} \\ & + \frac{\gamma^M}{24} C_1^M L^4 \sum_{ij} \int_{\Omega(\mathbf{x})} d^3\mathbf{x}' \mathcal{A}_{\alpha\zeta}\left(\mathbf{x}-\mathbf{x}'\right) \nabla'_\kappa \left\{ \left[\mathbf{x}_i - \mathbf{x}_j\right]_\beta \delta\left(\mathbf{x}-\mathbf{x}_i\right) \delta\left(\mathbf{x}'-\mathbf{x}_j\right) \left[\mathbf{p}_i - \mathbf{p}_j\right]_\kappa \left[\mathbf{v}_i - \mathbf{v}_j\right]_\zeta \right\} \\ & + \frac{\gamma^M}{24} C_1^M L^4 \sum_{ij} \int_{\Omega(\mathbf{x})} d^3\mathbf{x}' \mathcal{A}_{\alpha\zeta}\left(\mathbf{x}-\mathbf{x}'\right) \nabla'_\kappa \left\{ \left[\mathbf{x}_i - \mathbf{x}_j\right]_\beta \delta\left(\mathbf{x}-\mathbf{x}_i\right) \delta\left(\mathbf{x}'-\mathbf{x}_j\right) \left[\mathbf{p}_i - \mathbf{p}_j\right]_\kappa V_{||}\left(\mathbf{p}_i - \mathbf{p}_j\right)_\zeta \right\} \\ & + \mathcal{O}(L^4R^4),\end{aligned} \tag{73}$$

and

$$\begin{aligned}& \mathbf{\Sigma}^{M,2}_{\beta\alpha}(\mathbf{x}) = \\ & \frac{\gamma^M}{48} C_0^M L^4 \sum_{ij} \int_{\Omega(\mathbf{x})} d^3\mathbf{x}' \mathcal{A}_{\alpha\zeta}\left(\mathbf{x}-\mathbf{x}'\right) \nabla'^2_{\kappa\lambda} \left\{ \left[\mathbf{x}_i - \mathbf{x}_j\right]_\beta \delta\left(\mathbf{x}-\mathbf{x}_i\right) \delta\left(\mathbf{x}'-\mathbf{x}_j\right) \left[\mathbf{p}_i\mathbf{p}_i + \mathbf{p}_j\mathbf{p}_j\right]_{\kappa\lambda} \left[\mathbf{v}_i - \mathbf{v}_j\right]_\zeta \right\} \\ & + \frac{\gamma^M}{24} C_0^M L^4 \sum_{ij} \int_{\Omega(\mathbf{x})} d^3\mathbf{x}' \mathcal{A}_{\alpha\zeta}\left(\mathbf{x}-\mathbf{x}'\right) \nabla'^2_{\kappa\lambda} \left\{ \left[\mathbf{x}_i - \mathbf{x}_j\right]_\beta \delta\left(\mathbf{x}-\mathbf{x}_i\right) \delta\left(\mathbf{x}'-\mathbf{x}_j\right) \left[\mathbf{p}_i\mathbf{p}_i + \mathbf{p}_j\mathbf{p}_j\right]_{\kappa\lambda} \left[V_{||}\left(\mathbf{p}_i - \mathbf{p}_j\right)\right]_\zeta \right\} \\ & + \mathcal{O}(L^4R^4).\end{aligned} \tag{74}$$

Next, we execute the double sum leading to

$$\mathbf{\Sigma}^{M,0}_{\beta\alpha}(\mathbf{x}) = \frac{\gamma^M}{2} C_0^M \int_{\Omega(\mathbf{x})} d^3\mathbf{x}' \mathcal{A}_{\alpha\zeta}\left(\mathbf{x}-\mathbf{x}'\right) \rho\left(\mathbf{x}\right) \rho\left(\mathbf{x}'\right) \left[\mathbf{x}-\mathbf{x}'\right]_\beta \left[\mathbf{v}(\mathbf{x}) - \mathbf{v}(\mathbf{x}')\right]_\zeta + \mathcal{O}(L^4R^4) \tag{75}$$

$$\begin{aligned}\mathbf{\Sigma}^{M,1}_{\beta\alpha}(\mathbf{x}) = {} & \frac{\gamma^M}{24} C_0^M L^2 \int_{\Omega(\mathbf{x})} d^3\mathbf{x}' \mathcal{A}_{\alpha\zeta}\left(\mathbf{x}-\mathbf{x}'\right) \nabla'_\kappa \left\{ \left[\mathbf{x}-\mathbf{x}'\right]_\beta \rho\left(\mathbf{x}\right) \rho\left(\mathbf{x}'\right) \left[\mathcal{H}\left(\mathbf{x}\right) + \mathcal{H}\left(\mathbf{x}'\right)\right]_{\kappa\zeta} \right\} \\ & + \frac{\gamma^M}{24} C_1^M L^2 \sum_{ij} \int_{\Omega(\mathbf{x})} d^3\mathbf{x}' \mathcal{A}_{\alpha\zeta}\left(\mathbf{x}-\mathbf{x}'\right) \nabla'_\kappa \left\{ \left[\mathbf{x}-\mathbf{x}'\right]_\beta \rho\left(\mathbf{x}\right) \rho\left(\mathbf{x}'\right) \left[\mathcal{J}\left(\mathbf{x}\right) + \mathcal{J}\left(\mathbf{x}'\right) + \frac{2}{3} V_{||}\mathcal{I}\right]_{\kappa\zeta} \right\} \\ & + \mathcal{O}(L^4R^4),\end{aligned} \tag{76}$$

and

$$\begin{aligned}& \mathbf{\Sigma}^{M,2}_{\beta\alpha}(\mathbf{x}) = \\ & \frac{\gamma^M}{48} C_0^M L^2 \int_{\Omega(\mathbf{x})} d^3\mathbf{x}' \mathcal{A}_{\alpha\zeta}\left(\mathbf{x}-\mathbf{x}'\right) \nabla'^2_{\kappa\lambda} \left\{ \left[\mathbf{x}-\mathbf{x}'\right]_\beta \rho\left(\mathbf{x}\right) \rho\left(\mathbf{x}'\right) \left[\mathcal{K}\left(\mathbf{x}\right) - \mathcal{K}\left(\mathbf{x}'\right) + \frac{2}{3}\mathcal{I}\left(\mathbf{v}\left(\mathbf{x}\right) - \mathbf{v}\left(\mathbf{x}'\right)\right)\right]_{\kappa\lambda\zeta} \right\} \\ & + \mathcal{O}(L^4R^4).\end{aligned} \tag{77}$$

We next expand $\mathbf{x}'$ around $\mathbf{x}$. Thus for instance $\mathbf{v}(\mathbf{x}') = \mathbf{v}(\mathbf{x} + (\mathbf{x}' - \mathbf{x})) = \mathbf{v}(\mathbf{x}) + (\mathbf{x}' - \mathbf{x}) \cdot \nabla\mathbf{v}(\mathbf{x}) + \mathcal{O}\left((\mathbf{x}' - \mathbf{x})^2\right)$, yielding

$$\mathbf{\Sigma}^{M,0}_{\beta\alpha}(\mathbf{x}) = \frac{\gamma^M}{2} C_0^M \int_{\Omega(\mathbf{x})} d^3\mathbf{x}' \mathcal{A}_{\alpha\zeta}\left(\mathbf{x} - \mathbf{x}'\right) \rho^2\left(\mathbf{x}\right) \left[\mathbf{x} - \mathbf{x}'\right]_\beta \left[\mathbf{x} - \mathbf{x}'\right]_\kappa \nabla_\kappa \mathbf{v}_\zeta(\mathbf{x}) + \mathcal{O}(L^4R^4), \tag{78}$$

$$\begin{aligned}\mathbf{\Sigma}^{M,1}_{\beta\alpha}(\mathbf{x}) = &-\frac{\gamma^M}{12} C_0^M L^2 \int_{\Omega(\mathbf{x})} d^3\mathbf{x}' \mathcal{A}_{\alpha\zeta}\left(\mathbf{x} - \mathbf{x}'\right) \rho^2\left(\mathbf{x}\right) \mathcal{H}_{\beta\zeta}\left(\mathbf{x}\right) \\ &- \frac{\gamma^M}{12} C_1^M L^2 \int_{\Omega(\mathbf{x})} d^3\mathbf{x}' \mathcal{A}_{\alpha\zeta}\left(\mathbf{x} - \mathbf{x}'\right) \rho^2\left(\mathbf{x}\right) \left[\mathcal{J}_{\beta\zeta}\left(\mathbf{x}\right) + \frac{1}{3} V_{||} \delta_{\beta\zeta}\right] + \mathcal{O}(L^4R^4),\end{aligned} \tag{79}$$

and

$$\mathbf{\Sigma}^{M,2}_{\beta\alpha}(\mathbf{x}) = \frac{\gamma^M}{24} C_0^M L^2 \int_{\Omega(\mathbf{x})} d^3\mathbf{x}' \mathcal{A}_{\alpha\zeta}\left(\mathbf{x} - \mathbf{x}'\right) \rho^2\left(\mathbf{x}\right) \left[\nabla_\kappa \mathcal{K}_{\kappa\beta\zeta}\left(\mathbf{x}\right) + \frac{2}{3} \nabla_\beta \mathbf{v}_\zeta(\mathbf{x})\right] + \mathcal{O}(L^4R^4). \tag{80}$$

The relevant volume integrals over the sphere $\Omega$ are provided by Eq. (59) as well as

$$\int_{\Omega(\mathbf{x})} d^3\mathbf{x}' \mathcal{A}_{\alpha\zeta}\left(\mathbf{x} - \mathbf{x}'\right) \left[\mathbf{x} - \mathbf{x}'\right]_\beta \left[\mathbf{x} - \mathbf{x}'\right]_\kappa = \frac{4\pi R^5}{75} \left(\delta_{\alpha\zeta}\delta_{\beta\kappa} + \delta_{\beta\alpha}\delta_{\zeta\kappa} + \delta_{\alpha\kappa}\delta_{\beta\zeta}\right). \tag{81}$$

Performing these integrals and taking into account that $\mathcal{K} = 0$ for the isotropic case, after summing the terms $\mathbf{\Sigma}^{M,0}_{\beta\alpha}, \mathbf{\Sigma}^{M,1}_{\beta\alpha}, \mathbf{\Sigma}^{M,2}_{\beta\alpha}$ we arrive at

$$\begin{aligned}\mathbf{\Sigma}^M = &\gamma^M C_0^M \frac{4\pi R^5}{150} \rho^2 \left[\mathrm{Tr}\left(\nabla\mathbf{v}\right) \mathcal{I} + \nabla\mathbf{v} + \left(\nabla\mathbf{v}\right)^T\right] - \gamma^M C_1^M \frac{L^2}{12} \frac{4\pi R^3}{9} \rho^2 \left[\mathcal{J} + \frac{1}{3} V_{||} \mathcal{I}\right] \\ &+ \gamma^M C_0^M \frac{L^2}{12} \frac{4\pi R^3}{9} \rho^2 \left[\frac{1}{3} \nabla\mathbf{v} - \mathcal{H}\right].\end{aligned} \tag{82}$$

Following the same steps, we find for the stress induced by passive crosslinkers

$$\begin{aligned}\mathbf{\Sigma}^X = &\gamma^X C_0^X \frac{4\pi R^5}{150} \rho^2 \left[\mathrm{Tr}\left(\nabla\mathbf{v}\right) \mathcal{I} + \nabla\mathbf{v} + \left(\nabla\mathbf{v}\right)^T\right] - \gamma^X C_1^X \frac{L^2}{12} \frac{4\pi R^3}{9} \rho^2 \mathcal{J} \\ &+ \gamma^X C_0^X \frac{L^2}{12} \frac{4\pi R^3}{9} \rho^2 \left[\frac{1}{3} \nabla\mathbf{v} - \mathcal{H}\right].\end{aligned} \tag{83}$$

We use the expressions for $\mathcal{J}$, $\mathcal{H}$ and $\mathcal{K}$ that follow from single filament force and torque balances, respectively and arrive at an expression for the material stress $\mathbf{\Sigma} = \mathbf{\Sigma}^X + \mathbf{\Sigma}^M$ which reads

$$\mathbf{\Sigma}_{\alpha\beta}(x) \ = \ \eta \xi_{\alpha\beta\kappa\lambda} \nabla_\kappa \mathbf{v}_\lambda(\mathbf{x}) - \gamma^M V_{||} \frac{4\pi R^3}{27} \rho^2 \frac{L^2}{12} \left(C_1^M - C_0^M \frac{\gamma^M C_1^M + \gamma^X C_1^X}{\gamma^M C_0^M + \gamma^X C_0^X}\right) \mathcal{I}_{\alpha\beta}, \tag{84}$$

where

$$\eta = \frac{4\pi R^3}{15} \rho^2 \left(\gamma^X C_0^X + \gamma^M C_0^M\right) \left(\frac{R^2}{5} + \frac{L^2}{54}\right) \tag{85}$$

and

$$\xi_{\alpha\beta\kappa\lambda} = \xi_s \left[\frac{1}{2} \left(\delta_{\alpha\kappa}\delta_{\beta\lambda} + \delta_{\alpha\lambda}\delta_{\beta\kappa}\right) - \delta_{\alpha\beta}\delta_{\kappa\lambda}\right] + \xi_b \delta_{\alpha\beta}\delta_{\kappa\lambda}, \tag{86}$$

where $\xi_s = 1$ and $\xi_b = \frac{3}{2}$, leading to the shear viscocity being $\eta_s = \eta$ and the bulk viscocity being $\eta_b = \frac{3}{2}\eta$.

#### 3.3 Nonuniqueness of microscocopic models reflects morphogenetic redundancy in the cell cortex

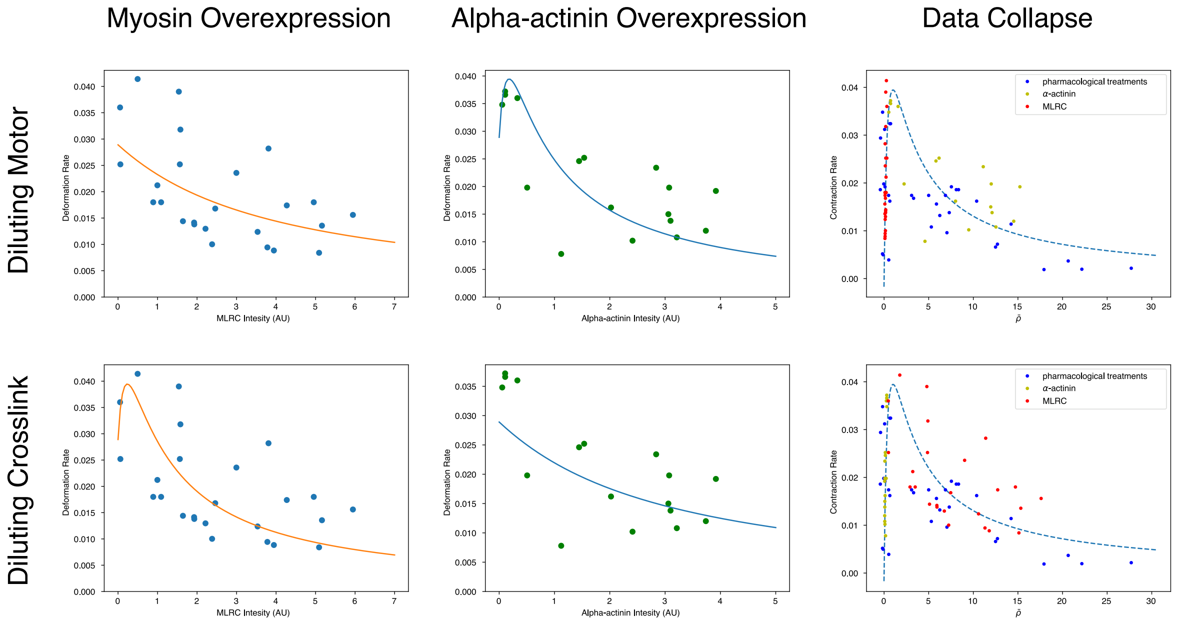

Figure S7: Fit comparison for models with where motor dilutes ($\mu^X - \mu^M = -1$) (top row) and for models where passive crosslinker dilutes ($\mu^X - \mu^M = 1$). Data in top row is identical to the one shown in Figs 5,6 of the main text.

In the main text the fits for Figs.(5,6) where presented under the choice that $\mu^M - \mu^X = -1$. This choice reflects microscopic models where the amount of motors per filament pair becomes smaller as more actin filaments become available, while the amount of crosslinks per actin filament pair stays constant. This can for instance be achieved if crosslink abundance is limited by binding site and motor abundance is limited by cutoplasmic concentration.

In this appendix we present the fit results for the opposite alternative $\mu^M - \mu^X = 1$, which could be achieved by a microscale physics in which the number of crosslinks per filament pair becomes smaller for higher actin densities and the amount of motor stays constant. As seen from the Fig.S7 This assumption produces fits which are similarly convincing as the choice presented in the main text in Figs (5,6). Thus, our large scale model can not uniquely decide between several possible microscale physics, see Fig. S7.

To clarify the issue, we summarize the simplest microscopic models, which differ by the end clustering behavior of motor and crosslink, and the values of $\mu^M - \mu^X$ respectively. We restrict ourselves to the simplest linear models, where either the per filament passive crosslinker or motor concentration scales inversely with actin density. Based on this choice of density scaling, as well as choices for the nonuniform localization of motors and crosslinkers a number of different models are possible. We begin by taking an agnostic approach and considering a large number of microscopic models listed in the table below. In all cases we assume that the motor moves toward the + end of the filament, i.e $V_{||} > 0$.

| Microscopic Model | Result |
|---|---|
| Uniform Motor & Crosslinker | $\Sigma_A = 0$, no active stress |
| End localizing Motor & Uniform Crosslinker<br>$C_0^M \sim \frac{C_0^M}{\rho}$ \| $C_1^M \sim \frac{C_1^M}{\rho}$ \| $C_0^X \sim C_0^X$ \| $C_1^X = 0$ | Strictly extensile |
| End localizing Motor & Uniform Crosslinker<br>$C_0^M \sim C_0^M$ \| $C_1^M \sim C_1^M$ \| $C_0^X \sim \frac{C_0^X}{\rho}$ \| $C_1^X = 0$ | Strictly extensile |
| Uniform Motor & End localizing Crosslinker at + end<br>$C_0^M \sim \frac{C_0^M}{\rho}$ \| $C_1^M = 0$ \| $C_0^X \sim C_0^X$ \| $C_1^X \sim C_1^X$ | **Plausible fits to data** |
| Uniform Motor & End localizing Crosslinker at + end<br>$C_0^M \sim C_0^M$ \| $C_1^M = 0$ \| $C_0^X \sim \frac{C_0^X}{\rho}$ \| $C_1^X \sim \frac{C_1^X}{\rho}$ | **Plausible fits to data** |
| Uniform Motor & End localizing Crosslinker at - end<br>$C_0^M \sim \frac{C_0^M}{\rho}$ \| $C_1^M = 0$ \| $C_0^X \sim C_0^X$ \| $C_1^X \sim -C_1^X$ | Strictly extensile |
| Uniform Motor & End localizing Crosslinker at - end<br>$C_0^M \sim C_0^M$ \| $C_1^M = 0$ \| $C_0^X \sim \frac{C_0^X}{\rho}$ \| $C_1^X \sim -\frac{C_1^X}{\rho}$ | Strictly extensile |

A unifying theme of the models consistent with the experimental measurements is that they break the filament-scale sliding asymmetry by having either an asymmetric driving force (through nonuniform motor localization) or an an asymmetric frictional force (through nonuniform crosslinker localization).

### 4 Extended Data Figures and Supplementary Videos

#### 4.1 Extended Data Figures

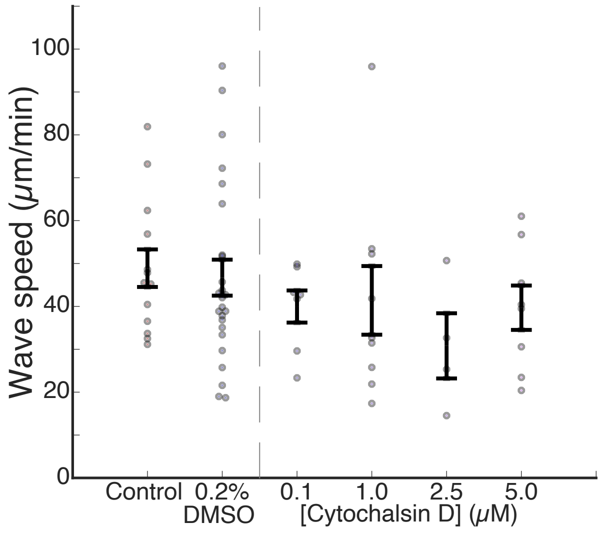

Extended Data Figure 1: SCW propagation speed does not change significantly under Cytochalasin D treatment. Errorbars: mean ± s.e.m. for each treatment condition.

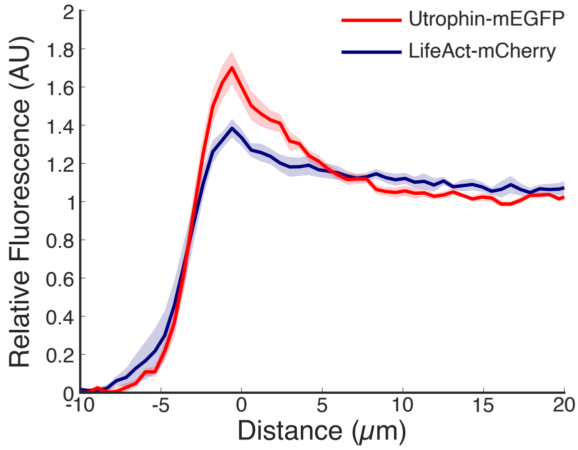

Extended Data Figure 2: Comparison between the fluorescence profiles when actin is visualized with either Utrophin-mEGFP or LifeAct-mCherry. For both Utrophin and LifeAct, fluorescence is peaked at the cortex and decays towards the oocyte's interior.

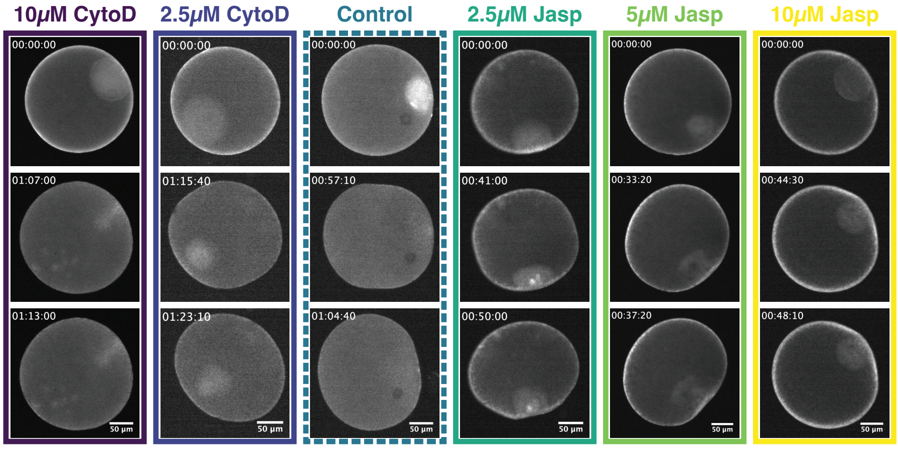

Extended Data Figure 3: Example images of LifeAct-mCherry fluorescence for cytochalsin D or jasplakinolide treated oocytes. Note that due to variable LifeAct-mCherry expression levels between oocytes, intensity values between oocytes cannot be directly compared.

#### 4.2 Supplementary Video Captions

**Supplementary Video 1: Surface contraction wave dynamics** Example of oocyte surface contraction wave. Time in hours: minutes: seconds. Colored circle surrounding oocyte indicates the local deformation rate. For colorbar, see Fig. 1c.

**Supplementary Video 2: LifeAct-mCherry as a proxy for F-actin** Example surface contraction wave of a control oocyte expressing LifeAct-mCherry. Time in hours: minutes: seconds.

**Supplementary Video 3: $\alpha$-actinin overexpression** Example surface contraction wave of an oocyte overexpressing $\alpha$-actinin-mEGFP. Time in hours: minutes: seconds.

**Supplementary Video 4: Myosin Regulatory Light Chain (MRLC) overexpression** Example surface contraction wave of an oocyte overexpressing MRLC-mEGFP. Time in hours: minutes: seconds.

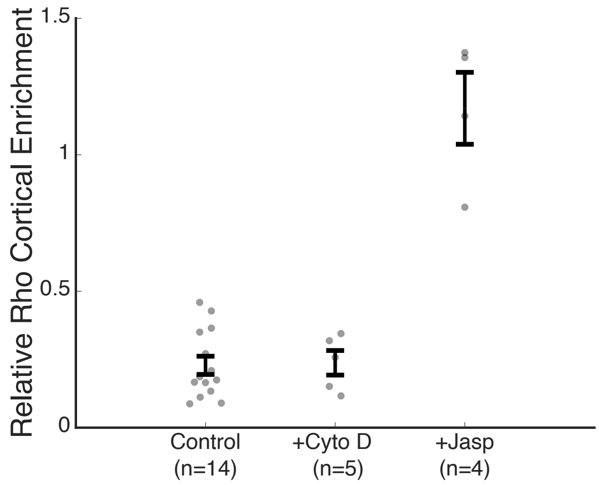

Extended Data Figure 4: Relative cortical enrichment of active Rho measured using rGBD-GFP for oocytes treated with 10$\mu$M cytochalasin d, 10$\mu$M jasplakinolide, or DMSO alone.

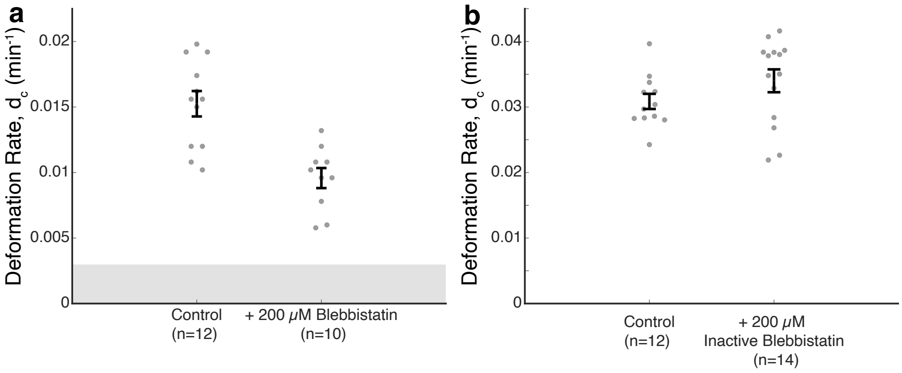

Extended Data Figure 5: (a) Partial myosin inhibition through blebbistatin treatment decreases the characteristic deformation rate. Data collected using brightfield microscope. (b) Treatment with an inactive blebbistatin enantiomer does not significantly change the deformation rate. Data collected using spinning disk microscope. Errorbars: mean ± s.e.m.

### References

1. Tan, T. H. *et al.* Topological turbulence in the membrane of a living cell. *Nature Physics* **16,** 657–662 (2020).
2. Mori, M. *et al.* An Arp2/3 Nucleated F-Actin Shell Fragments Nuclear Membranes at Nuclear Envelope Breakdown in Starfish Oocytes. *Current Biology* **24,** 1421–1428 (2014).
3. Bischof, J. *et al.* A cdk1 gradient guides surface contraction waves in oocytes. *Nature Communications* **8,** 1–10 (2017).
4. Edelstein, A. D. *et al.* Advanced methods of microscope control using $\mu$Manager software. *Journal of biological methods* **1** (2014).
5. Klughammer, N. *et al.* Cytoplasmic flows in starfish oocytes are fully determined by cortical contractions. *PLoS Computational Biology* **14,** e1006588 (2018).
6. Mietke, A., Jülicher, F. & Sbalzarini, I. F. Self-organized shape dynamics of active surfaces. *Proc Natl Acad Sci U S A* **116,** 29–34 (Jan. 2019).

7. Bächer, C., Khoromskaia, D., Salbreux, G. & Gekle, S. A Three-Dimensional Numerical Model of an Active Cell Cortex in the Viscous Limit. *Frontiers in Physics* **9,** 753230 (2021).
8. Borja da Rocha, H., Bleyer, J. & Turlier, H. A viscous active shell theory of the cell cortex. *Journal of the Mechanics and Physics of Solids* **164,** 104876 (2022).
9. Wittwer, L. D. & Aland, S. A computational model of self-organized shape dynamics of active surfaces in fluids. *Journal of Computational Physics: X* **17,** 100126 (2023).
10. Fürthauer, S., Needleman, D. J. & Shelley, M. J. A design framework for actively crosslinked filament networks. *New Journal of Physics* **23,** 013012 (2021).
11. Mayer, M., Depken M.and Bois, J., Jülicher, F. & Grill, S. W. Anisotropies in cortical tension reveal the physical basis of polarizing cortical flows. *Nature* **467,** 617â€"621 (2010).
12. Hou, T. Y., Lowengrub, J. S. & Shelley, M. J. Removing the stiffness from interfacial flows with surface tension. *Journal of Computational Physics* **114,** 312–338 (1994).
